# Three STEPs forward: A trio of unexpected structures of PTPN5

**DOI:** 10.1101/2024.11.20.624168

**Authors:** Liliana Guerrero, Ali Ebrahim, Blake T. Riley, Sean H. Kim, Anthony C. Bishop, Jiaqian Wu, Ye Na Han, Lutz Tautz, Daniel A. Keedy

## Abstract

Protein tyrosine phosphatases (PTPs) play pivotal roles in myriad cellular processes by counteracting protein tyrosine kinases. Striatal-enriched protein tyrosine phosphatase (STEP, PTPN5) regulates synaptic function and neuronal plasticity in the brain and is a therapeutic target for several neurological disorders. Here, we present three new crystal structures of STEP, each with unexpected features. These include high-resolution conformational heterogeneity at multiple sites, and a highly coordinated citrate molecule in the active site, a previously unseen conformational change at an allosteric site, an intramolecular disulfide bond that was characterized biochemically but had never been visualized structurally, and two serendipitous covalent ligand binding events at surface-exposed cysteines that are nearly or entirely unique to STEP among human PTPs. Together, our results offer new views of the conformational landscape of STEP that may inform structure-based design of allosteric small molecules to specifically inhibit this biomedically important enzyme.

## Introduction

Protein tyrosine phosphatases (PTPs) counteract protein tyrosine kinases to regulate many cellular processes. They do so by dephosphorylating post-translationally modified phosphotyrosine (pTyr) moieties in a variety of specific substrate proteins in human cells. Striatal-enriched protein tyrosine phosphatase (STEP, i.e. PTPN5) is a PTP found primarily in striatal neurons ^1^. A key substrate of STEP is the N-methyl-_D_-aspartate (NMDA) receptor, which is a glutamate neurotransmitter receptor that plays key roles in neuroplasticity and learning ^2^. Due to its important role in the brain, STEP is a validated therapeutic target for Fragile X syndrome ^3^, Parkinson’s disease ^4^, and Alzheimer’s disease ^5,6^, as supported by mouse knockout models ^3,7–9^.

Several small-molecule inhibitors targeting the active site of STEP have been reported, with associated crystal structures ^10,11^. However, because the active site is highly charged and highly conserved among PTPs, active-site inhibitors generally suffer from bioavailability and selectivity limitations, respectively ^12–14^. Unusually for PTPs, an allosteric small-molecule activator of STEP has also been reported, including crystal structures of two compound variants ^15^. However, these activators are relatively weak (EC_50_ 100–500 µM), and do not address the need for inhibitors of STEP. Altogether, no potent, specific small-molecule inhibitors of STEP have been introduced, highlighting the importance of identifying new ligandable allosteric sites in STEP’s 3-dimensional structure that could enable different strategies for drug design.

Despite this gap for small-molecule inhibition, STEP is known to exhibit complex natural regulatory mechanisms. For example, it is highly sensitive to oxidative stress, which can inactivate the enzyme through the formation of a disulfide bond between its catalytic cysteine and a “backdoor” cysteine ^16^. This reversible oxidation process serves as a key regulatory mechanism in STEP^17^ and other PTPs ^18–20^. Additionally, STEP’s enzymatic activity is regulated by various post-translational modifications such as phosphorylation and ubiquitination, which can influence STEP’s stability, localization, and interactions with other proteins ^13^. Together, these observations suggest that the structure of STEP may be tuned to respond to a variety of regulatory inputs, presenting potential opportunities for design of new allosteric modulators that exploit its inherent biophysical properties.

Here we report three serendipitous new crystal structures of STEP, all of which include unexpected features that provide complementary and unique insights into the potential allosteric druggability of this important protein. (1) First, we report the highest-resolution STEP structure to date by a considerable margin, which offers a detailed view of conformational heterogeneity throughout the protein. This structure also features a previously unseen ordered citrate molecule bound in a highly coordinated fashion in the active site region. (2) Second, in a structure derived from unintended crystal dehydration, we observe a distinctive conformational change in the allosteric S loop, part of the binding site for the previously reported small-molecule allosteric activator ^15^. This change opens up the site, providing additional binding pocket volume that may guide future structure-based design of optimized allosteric activators or inhibitors. Additionally, this structure reveals an intramolecular disulfide bond that has not been previously visualized, shining atomistic light on a recognized regulatory mechanism of STEP. (3) Third, we report a structure of STEP in complex with a known active-site inhibitor of other PTPs ^21,22^: the <u>co</u>valent ligand 2-[4-(2-<u>br</u>omoacetyl)phenoxy]-<u>a</u>cetic acid (hereafter referred to as “CoBrA”). Despite this, we unequivocally demonstrate covalent binding at two surface-exposed cysteines in STEP, both distal from the active site. As one of these cysteines is almost unique and the other is entirely unique to STEP among human PTPs, the binding events in this structure offer novel footholds for the design of STEP-specific allosteric small-molecule modulators.

Altogether, the trifecta of new structures we report here provides unique opportunities for rational structure-based drug discovery of allosteric small-molecule modulators for STEP, and for augmenting our understanding of the endogenous regulatory mechanisms of this biomedically important enzyme in the human brain.

## Results

### Overview of new structures

Here we report three unexpected X-ray crystal structures of STEP. We used the STEP catalytic domain for our experiments, as in previous structures ^23^. The X-ray data reduction and model refinement statistics for our datasets are favorable (**Table 1**). All the resolutions are 1.75 Å or greater, including 1.27 Å for one dataset, which is the highest resolution for any STEP crystal structure (**Fig. 2a**), and among the highest for any PTP.

**Table 1:** Crystallographic statistics. Overall statistics given first (statistics for highest-resolution bin in parentheses).

|  |  |  |  |
| --- | --- | --- | --- |
| PDB ID | 9EEX | 9EEY | 9EEZ |
| Structure | Citrate-bound STEP | Dehydrated STEP | Covalently bound STEP |
| Resolution (Å) | 58.03 – 1.27 | 63.62 – 1.75 | 64.20 – 1.60 |
| Data collection temperature (K) | 100 | 100 | 100 |
| Completeness (%) | 99.9 (99.0) | 99.8 (98.7) | 99.9 (99.0) |
| Multiplicity | 12.5 (12.2) | 12.9 (11.2) | 12.4 (12.6) |
| I/sigma(I) | 9.2 (1.1) | 7.2 (0.3) | 13.2 (0.8) |
| R<sub>merge</sub>(I) | 0.157 (4.132) | 0.157 (4.668) | 0.084 (2.800) |
| R<sub>meas</sub>(I) | 0.163 (4.313) | 0.164 (4.890) | 0.088 (2.922) |
| R<sub>pim</sub>(I) | 0.046 (1.217) | 0.045 (1.443) | 0.025 (0.820) |
| CC<sub>1/2</sub> | 0.998 (0.364) | 0.997 (0.320) | 0.999 (0.406) |
| Wilson B-factor (Å<sup>2</sup>) | 13.89 | 36.91 | 27.63 |
| Total observations | 1172163 (55967) | 460854 (19279) | 590840 (28520) |
| Unique observations | 93644 (4593) | 35607 (1726) | 47763 (2270) |
| Space group | P 21 21 21 | P 21 21 21 | P 21 21 21 |
| Unit cell dimensions (Å, Å, Å, °, °, °) | 40.105, 64.154, 136.043, 90, 90, 90 | 39.673, 63.621, 135.800, 90, 90, 90 | 40.144, 64.194, 137.182, 90, 90, 90 |
| Solvent content (%) | 54.31 | 53.36 | 54.75 |
| R<sub>work</sub> | 0.131 | 0.199 | 0.202 |
| R<sub>free</sub> | 0.153 | 0.238 | 0.235 |
| RMS bonds (Å) | 0.006 | 0.010 | 0.011 |
| RMS angles (°) | 0.86 | 0.87 | 0.99 |
| Ramachandran outliers (%) | 0.36 | 0.38 | 0.38 |
| Ramachandran favored (%) | 96.43 | 93.61 | 95.83 |
| Clashscore | 2.79 | 0.45 | 2.01 |
| MolProbity score | 1.30 | 1.08 | 1.26 |

The overall protein fold in these structures is similar to previous structures of STEP. However, our new structures reveal several interesting and unexpected features, including (i) a citrate molecule bound to the active site of STEP at high resolution, (ii) a shifted loop conformation that opens up an allosteric activator pocket, (iii) an intramolecular disulfide bond involving the catalytic cysteine, and (iv) a covalent ligand bound at surface sites distal from the active site (**Fig. 1**). The remainder of the paper explores these structural observations in detail.

**Figure 1.**
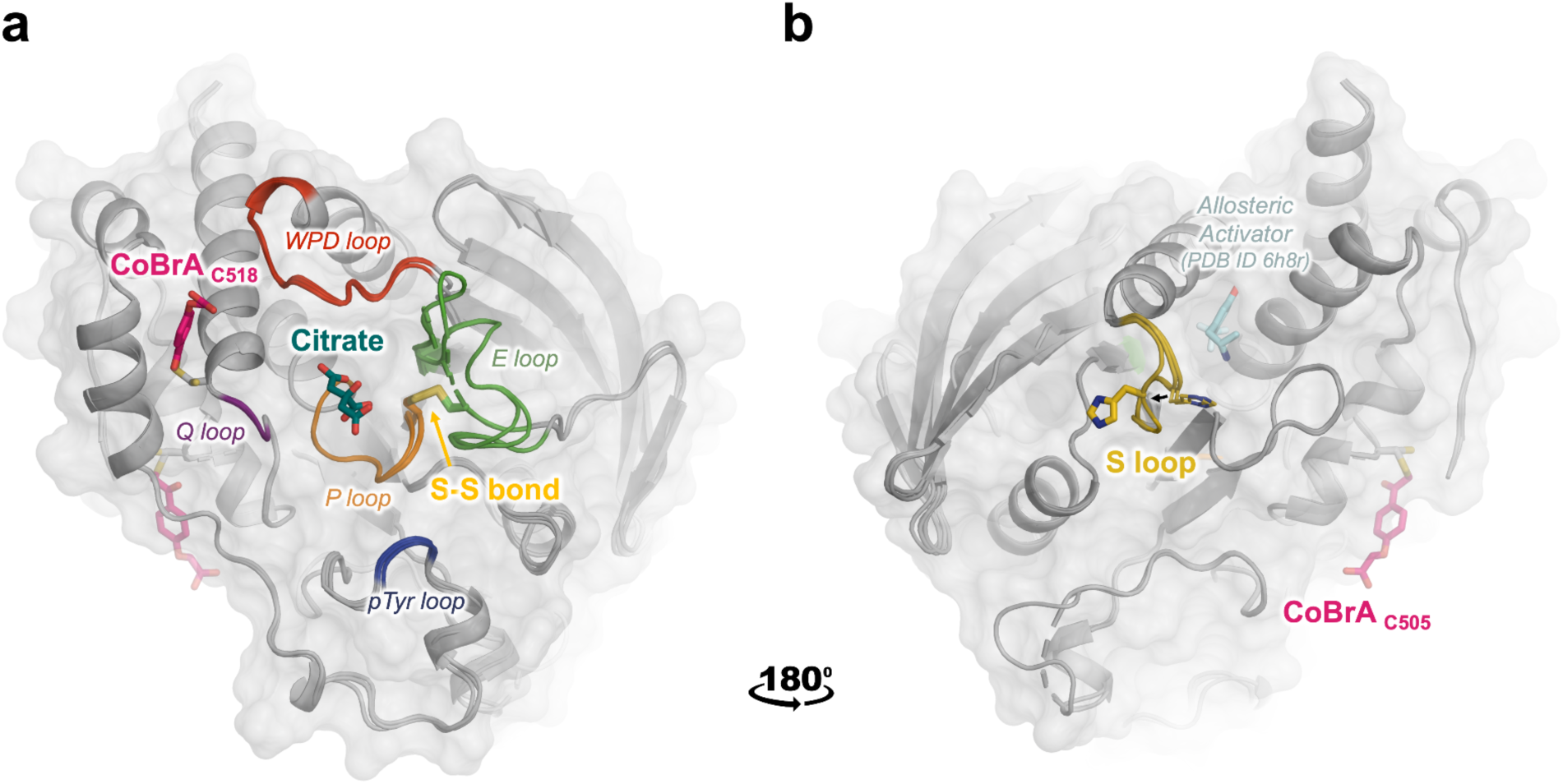
Overview of three unexpected new structures of STEP. Overlay of three new crystal structures of STEP, with conformational changes and binding events at both known and novel ligand-binding sites. a) Citrate is non-covalently bound in the active site (dark green sticks, center) amidst nearby active-site loops. The “CoBrA” covalent ligand is bound at Cys518 (pink sticks, left). Also shown is the intramolecular disulfide bond between the catalytic Cys472 and the nearby “backdoor” Cys384, denoted as S-S bond (gold sticks). b) View rotated by 180°. The distal S loop (yellow) undergoes a conformational shift in the pocket that was shown to bind small-molecule allosteric activators in prior structures ^15^ (pale cyan sticks); His464 shifts away from the position of the allosteric activator. CoBrA is also covalently bound at Cys505 (pink sticks, right).

**Figure 2.**
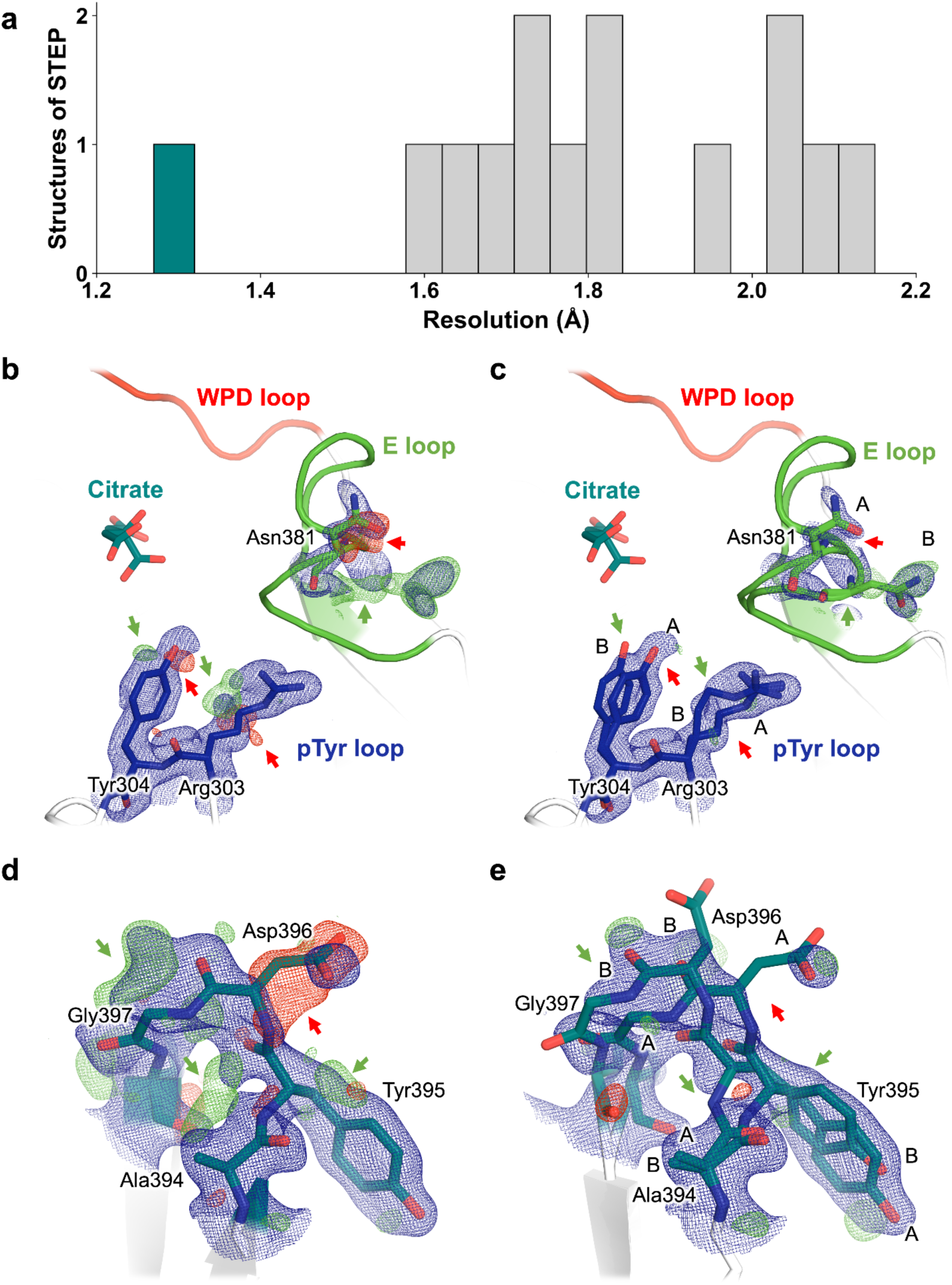
Protein conformational heterogeneity revealed at high resolution. **a)** Histogram of resolution for all previous structures of STEP from the PDB, plus our new high-resolution citrate-bound structure (1.27 Å) in dark green. The other structures reported in this work are also included in the histogram. **b,c)** Single-conformer model for high-resolution dataset with 2Fo-Fc electron density (1 σ, blue) for several sites in STEP. The Fo-Fc density (+/- 3 σ, green/red) suggests missing, unmodeled alternate conformations (arrows). **d,e)** Final multiconformer model with 2Fo-Fc electron density (1 σ, blue) for the same sites. The reduced Fo-Fc density (+/- 3 σ, green/red), indicated by arrows, demonstrates that the multiconformer model is an improved representation for these sites.

### Bound citrate at high resolution

As noted above, we report the highest-resolution X-ray crystallography dataset of STEP to date, at 1.27 Å. This readily surpasses previous STEP structures, which have resolutions ranging from 1.66–2.15 Å (**Fig. 2a**), and is within the top 1–2% of all PTP family structures in the Protein Data Bank (PDB) ^24,25^. One benefit of a very high-resolution crystal structure is that we are able to observe clear evidence for alternate conformations in numerous locations. For example, Arg303–Tyr304 in the pTyr binding loop (also known as the substrate binding loop or substrate recognition loop) adopt alternate side-chain conformations, and the spatially adjacent Asn381 and neighboring residues in the flexible active-site E loop exhibit coupled side-chain and backbone conformational heterogeneity (**Fig. 2b-c**). Distal from the active site, several residues in a β-hairpin also display correlated backbone displacements (**Fig. 2d-e**).

In previous crystal structures, STEP has only been observed with an atypically “super-open” WPD loop conformation that is incompatible with the known catalytic mechanism, as opposed to the typical open and closed states seen in most PTPs (**Fig. 3d**). This may be because, in all but one previous structure of STEP, a competitive inhibitor or an ordered sulfate molecule from the crystallization solution binds in a subsite of the active-site pocket that would block closure of the WPD loop ^23^. In the remaining prior structure, a pTyr substrate is bound to the main catalytic subsite; this structure accordingly shows some global signatures of an active-like state ^23^, but the WPD loop remains atypically open, possibly due in part to the C472S mutation of the catalytic cysteine that was necessary to capture this complex. The atypically open conformation of the WPD loop in STEP is also thought to be stabilized by other structural features ^26,27^.

**Figure 3.**
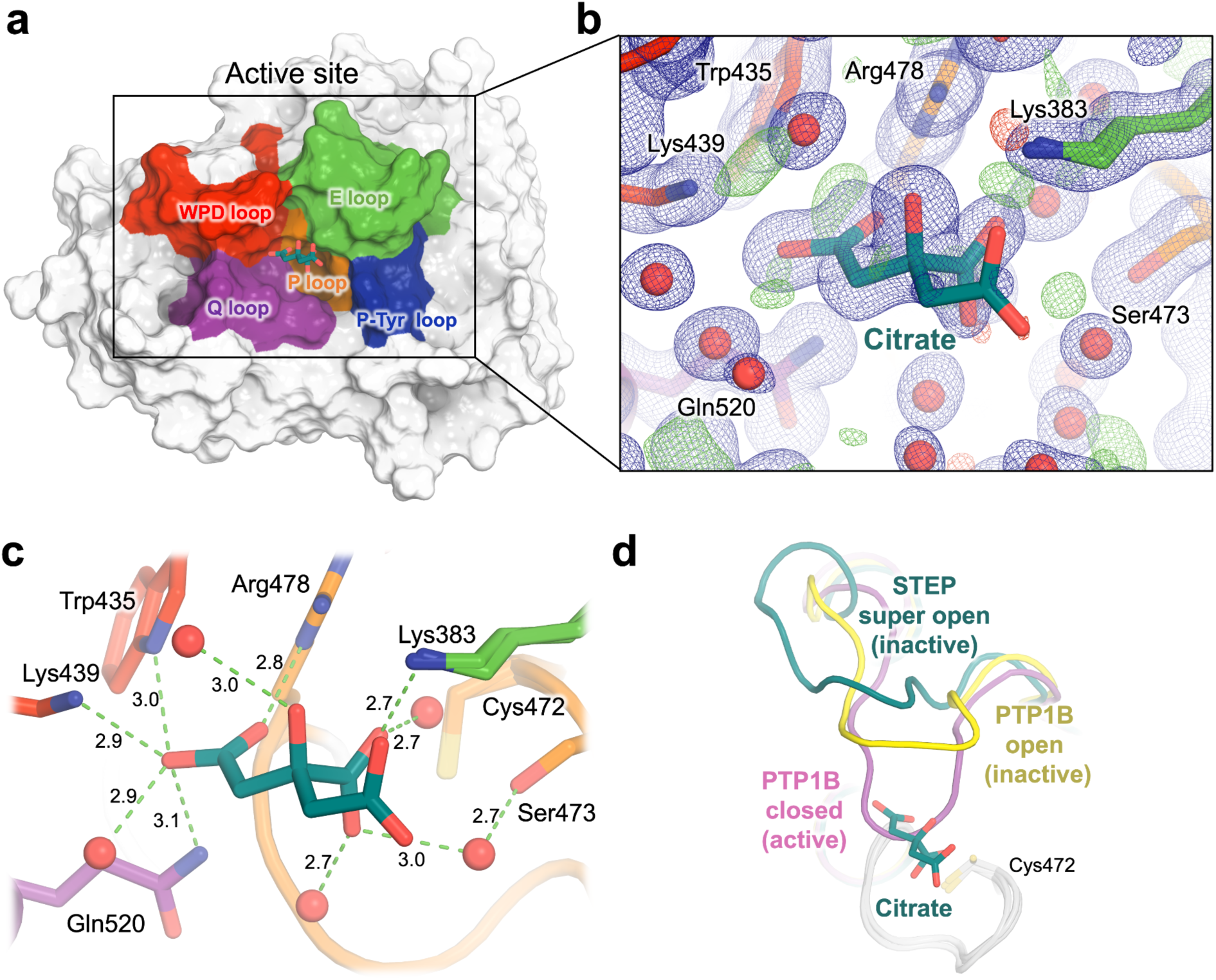
Crystal structure of STEP bound to highly coordinated citrate. **a)** Overview of STEP with bound citrate in the active site, and key catalytic loops in different colors. **b)** Zoom-in to citrate, showing our refined structural model with strong support from 2Fo-Fc (1 σ in blue) and Fo-Fc (+/- 3 σ in green/red) electron density. **c)** Interactions of citrate with surrounding amino acids in the active site pocket, including several H-bonds (dotted green lines) and no steric clashes. Distances are given in Å. The catalytic Cys472 is nearby, although it does not directly interact with the citrate. **d)** In our structure (dark green), the WPD loop is in an atypically “super-open” state, as with all previous STEP structures. However, the bound citrate is mutually exclusive with the canonical closed state of the WPD loop from PTP1B (PDB ID 1pty, pink), as distinct from the open state (PDB ID 1t49, yellow).

In an attempt to alter the WPD loop state, we crystallized STEP in different conditions with citrate instead of sulfate (see Methods), resulting in the high-resolution dataset presented here. Modulating the WPD loop conformation was unsuccessful; however, unexpectedly, the high-resolution map reveals clear 2Fo-Fc electron density on the periphery of the active site pocket, which we interpret as a non-covalently bound citrate molecule (**Fig. 3a-b**). There are no crystal lattice contacts to the citrate or any nearby protein or loop residues, arguing against the notion that the clear presence of this molecule in our structure is an artifact of the crystal environment.

The citrate is well-coordinated by numerous protein side chains and ordered water molecules in the active site environment, forming a network of 10 hydrogen bonds (**Fig. 3c**). These interactions include H-bonds to the conserved Gln520 of the conserved catalytic Q loop ^28^ and to a well-ordered water molecule (B-factor: 23.60 Å^2^) ^29^ that is also coordinated by Gln520; an H-bond to Lys383 of the flexible E loop flanking the active site; and H-bonds to the essential and conserved Arg478 of the catalytic P loop. The citrate is also linked to Ser473 through a water bridge. Furthermore, the citrate forms an H-bond with the poorly conserved Lys439 region of the WPD loop in STEP, which is only present in STEP and its homolog PTPRR ^23,30^ (**Fig. 3c)**.

Citrate binding occurs in a location that is mutually exclusive with the putative closed, or “active”, conformation of the WPD loop (**Fig. 3d**). To test the functional relevance of this observed interaction between STEP and citrate in solution, we performed an *in vitro* enzyme activity assay. In this assay, buffer concentration was chosen to ensure that assay pH remained stable with increasing amounts of citrate. The results showed a weak reduction of STEP enzyme activity by citrate with an IC_50_ of 6.4 mM (**Fig. S1**), consistent with our structural findings (**Fig. 3**).

Although not previously seen in crystallographic structures of STEP, citrate has been linked to PTP binding and activity in the PDB and the literature. For instance, an ordered citrate, presumably from the crystallization solution, was recently observed in the active site of a crystal structure of SHP2 bound to a monobody, albeit in a different pose than in our structure (PDB ID 7tvj) ^31^ (**Fig. S2**). In addition, iron-citrate complexes were shown to competitively inhibit the enzymatic activity of SHP1 ^32^. Thus, there is precedent for various forms of citrate to interact with PTP active sites.

### Allosteric loop shift and intramolecular disulfide bond

The second surprising structure of STEP that we report derives from a crystal subjected to slow dehydration for ∼6 months prior to crystal harvesting (see Methods). This long incubation of the crystallization drops has resulted in several intriguing changes to the structure of STEP.

Relative to the other datasets from this study, the unit cell dimensions of this dehydrated crystal structure decrease by ∼1% (**Table 1**), with the resulting unit cell volume decreasing by 3.0% and 2.1% versus the citrate-bound and covalently bound structures, respectively. These changes are similar in magnitude to the decrease in unit cell volume of 3.4% upon crystal cryocooling, discerned in a prior study by comparing room-temperature (RT) to cryogenic (cryo) crystal structures for many proteins ^33^. These changes are also similar to the decreases in unit cell volume of 3.9% for ambient-pressure cryocooling and 2.1% for high-pressure cryocooling that we reported previously for this crystal form of STEP ^23^. Dehydration, therefore, appears to have had a similarly large effect on STEP in the crystal environment as do other important biophysical perturbations.

Perhaps as a result of this apparent dehydration, we observe clear evidence for a significant conformational change of the non-conserved ^15,34^, surface-exposed S loop (residues 462–467) (**Fig. 4a**), located ∼20 Å from the active site. It is not immediately obvious what other structural changes favor this loop conformational change, but subtle alterations of the crystal lattice contacts involving Pro463–His464 may play a role.

**Figure 4.**
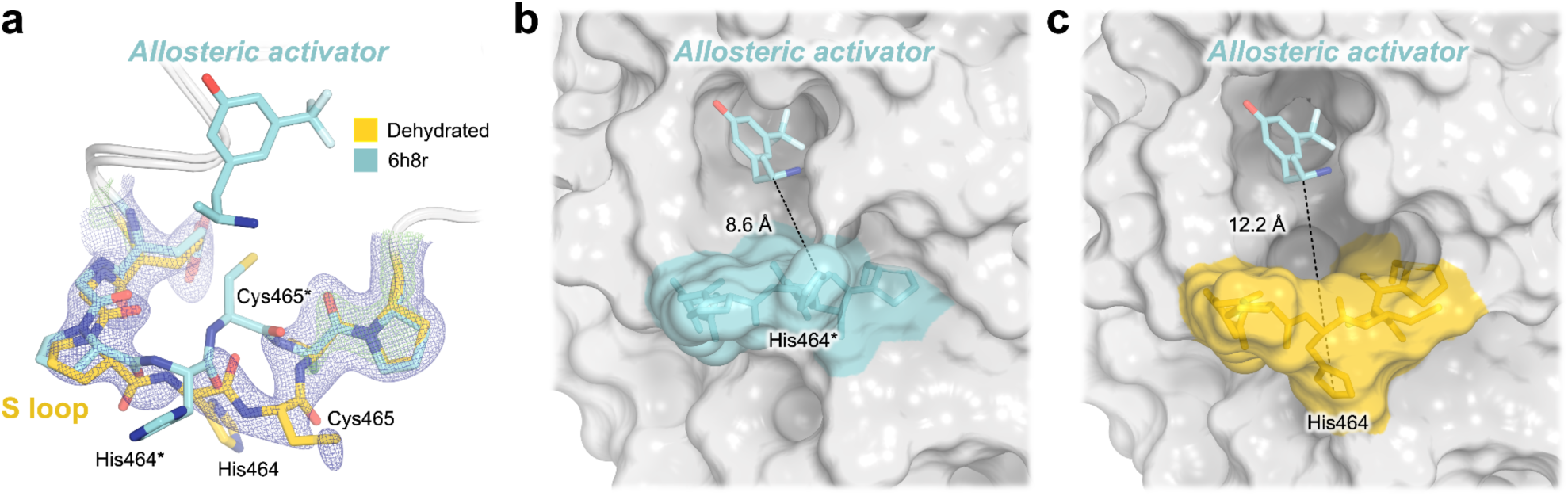
Crystal structure shows unanticipated rearrangement of allosteric S loop. **a)** 2Fo-Fc electron density for a distinct conformation of the S loop in our new structure of STEP. This conformation differs from that seen in a previous structure bound to an allosteric small-molecule activator in the adjacent pocket (PDB ID 6h8r) ^15^. **b-c)** The conformational change of the loop in our structure opens up the activator binding site pocket, as measured by an increased distance between a central residue (His464) in the S loop and the allosteric activator from 6h8r. Asterisks (*) indicate our renumbering of the non-standard residue numbering in 6h8r (see Methods).

Notably, the S loop forms a key part of the previously reported allosteric site in STEP, interrogated in the past with two variants of a small-molecule allosteric activator ^15^. To date, these compounds constitute the only allosteric modulators of STEP, although their potencies are limited, with 500 µM compound required for >50% activation of human STEP ^15^. In previous crystal structures of human STEP and mouse STEP bound to two different activator variants, the S loop did not deviate substantially from its position relative to apo structures despite binding of the ligand nearby. By contrast, the new conformation of the S loop in our dehydrated structure is distinct from that seen in all previous structures of STEP (**Fig. S3**). Unlike the previous structure of mouse STEP with an allosteric activator in which the loop “closes” slightly into the pocket (PDB ID 6h8s), our structure shows the loop becoming more “open” (**Fig. 4b-c**). Strikingly, this shift causes the allosteric pocket to double in volume, from 83.4 Å^3^ in the activator-bound structure of human STEP (PDB ID 6h8r) to 166.9 Å^3^ in our structure.

A second surprising feature we observe from this dehydrated dataset is electron density consistent with an intramolecular disulfide bond from the catalytic cysteine (Cys472) to the nearby “backdoor” cysteine (Cys384) from the active-site E loop (**Fig. 5**). It is possible that an oxidizing environment arose during the long incubation and slow dehydration of the crystal, contributing to formation of this disulfide bond. Our observation of a disulfide is consistent with past biochemical experiments showing that STEP forms a reversible intramolecular disulfide bond between these cysteines in response to oxidation, a behavior which was not observed in another KIM-type PTP, HePTP ^16^.

**Figure 5.**
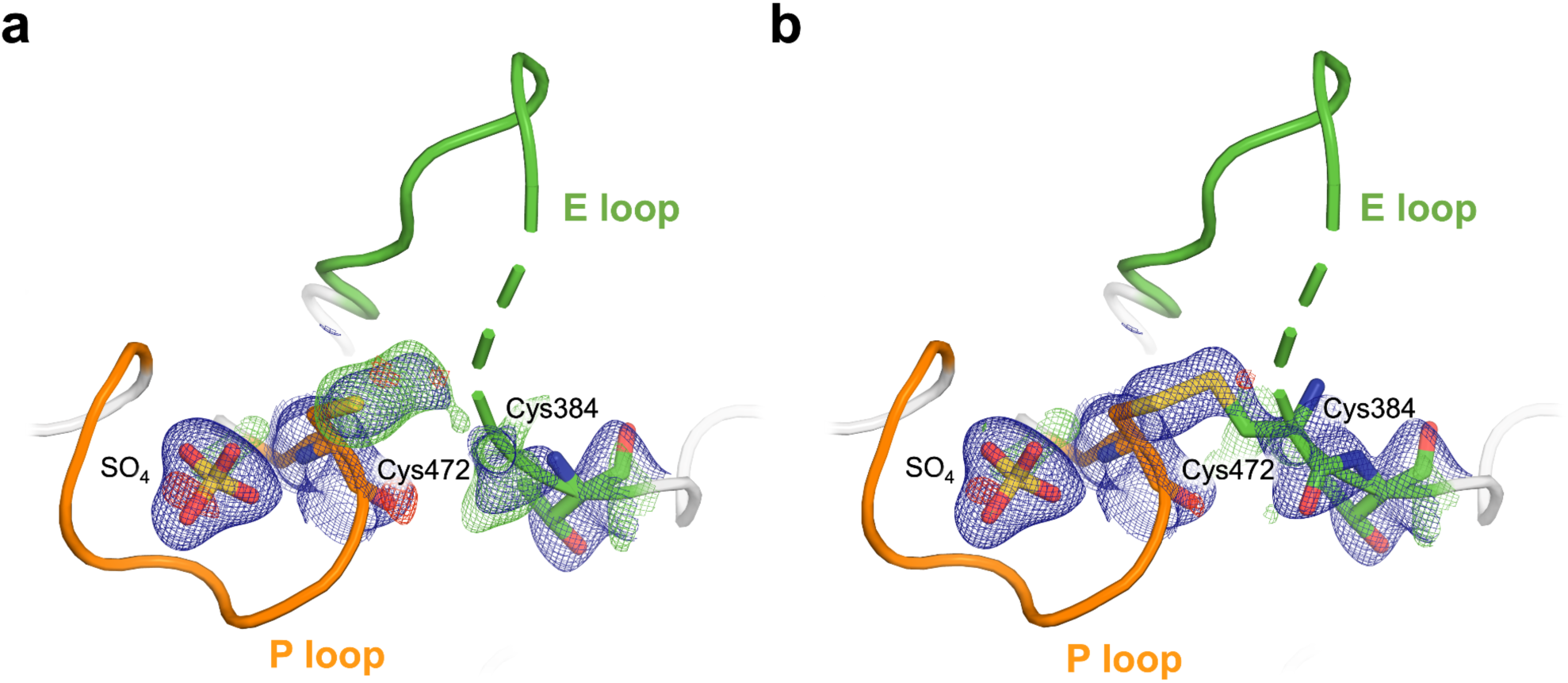
Intramolecular disulfide bond to backdoor cysteine in dehydrated STEP structure. **a)** Omit electron density (2.7 σ in green; Cys384 and Sγ atom of Cys472 omitted), showing unbiased evidence of an intramolecular disulfide bond between the catalytic Cys472 and the nearby “backdoor” Cys384 from the E loop. **b)** 2Fo-Fc (1.0 σ in blue) electron density for our refined structure with the disulfide bond modeled, showing good fit to density. The portion of the E loop immediately N-terminal to Cys384 was observed to be disordered upon disulfide formation (indicated by dashed cartoon representation).

As this disulfide has not previously been modeled in any structure of STEP, we assessed the degree to which our new dataset is unique for this disulfide region. To test whether a disulfide may have been present but unmodeled in the previous structures, we examined electron density within the catalytic pocket from several previous structures of STEP, as well as the three structures in this study. For each, we modeled a putative disulfide bond, re-refined against the corresponding structure factors, and then examined the resulting electron density maps. For the previously deposited structures of STEP, we see little to no density consistent with the disulfide (**Fig. S5a-f**), with the same being true of our high-resolution, citrate-bound structure (**Fig. S5g**). By contrast, the electron density is clear and irrefutable for the disulfide in our dehydrated dataset (**Fig. S5h**), as well as the covalently bound dataset from this study (**Fig. S5i**; see next section).

During initial refinement of our new structures, negative density peaks near the sulfur atoms participating in the disulfide bonds were observed (**Fig. S6a,c**). This negative density suggests radiation-induced disruption of the disulfide bond, an event commonly observed in X-ray crystallography ^35–37^. We addressed this by modeling the sulfurs with partial occupancies, which resolved the negative difference density (**Fig. S6b,d**).

Furthermore, in structural modeling during refinement, we noticed that the E loop region immediately preceding the backdoor Cys384 (typically residues Ile374–Lys383) is disordered based on 2Fo-Fc electron density or lack thereof in both the dehydrated and covalently bound STEP structures (**Fig. 5**, dashed cartoon). This region, along with the loop preceding the pTyr loop (residues Lys291–Arg300) and the loop formed by residues Thr318–Leu326, are in close contact within the crystal lattice in this crystal form (**Fig. S7**). Due to the lack of continuous electron density in these regions, these three loops are challenging to model. We hypothesize that formation of the Cys472-Cys384 disulfide bond may trigger movement in the E loop that then affects adjacent loops in the crystal. Interestingly, however, oxidation of the catalytic cysteine in the STEP homolog PTP1B has also been associated with a conformational change in the loop region corresponding to STEP residues Leu317–Tyr329 ^38^, suggesting potential allosteric regulation within the PTP domain itself that is linked to the oxidation state of the catalytic cysteine.

In line with our observations of potential oxidation at these two active-site cysteines in our dehydrated dataset, we also observe extended electron density at two distal cysteines, Cys505 and Cys518, which may suggest covalent modifications such as the addition of oxygen atoms (**Fig. S8b,f**).

Intriguingly, several previous structures of oxidized STEP modeled covalent modifications that include additional atoms at Cys518; however, these models are not consistent with our electron density maps (**Fig. S8h**). We have, therefore, left these regions unmodeled in our dehydrated structure (see Methods). Nevertheless, these observations indicate that Cys505 and Cys518 in STEP may be chemically reactive — a finding which is further exemplified by the third STEP structure reported here.

### Covalent ligand at non-catalytic cysteines

Previously, a series of covalent, photocleavable inhibitors targeting the catalytic cysteine of a small panel of PTPs were reported ^21,22^. These α-haloacetophenone derivative compounds were shown to inhibit multiple PTPs (albeit with different potencies), which is unsurprising given the generally conserved nature of the catalytic pocket within PTP family enzymes. As STEP was not among the PTPs tested in that work, we sought to test whether the <u>co</u>valent binder 2-[4-(2-<u>br</u>omoacetyl)phenoxy]-<u>a</u>cetic acid (here “CoBrA”) would also form a covalent adduct to the catalytic cysteine in STEP (Cys472), and thus bring about previously unseen conformational changes in STEP to provide insight into its mechanisms of catalysis and/or allosteric regulation.

Notably, we observed no evidence of covalent binding to the catalytic cysteine, Cys472, in our crystal structure. Although electron density is present in the active-site pocket, it is not continuous with the electron density for the side chain of Cys472 as would be expected for a covalent linkage (**Fig. S9a**). In this structure, we can attribute the density in the pocket to a sulfate that mimics the pTyr substrate, as seen previously in multiple STEP crystal structures ^23^. Attempting to model CoBrA at this site leads to a poor fit to the electron density (**Fig. S9b**), arguing against its presence in the active site.

To our surprise, we do observe strong 2Fo-Fc and Fo-Fc electron density supporting covalent binding of CoBrA at the two distal cysteines mentioned previously (**Fig. S8**), Cys505 and Cys518, which are

∼21 Å and ∼7 Å from the active site, respectively. 2Fo-Fc and unbiased omit electron density maps are continuous for CoBrA, while the side-chain conformations at these two cysteines are also consistent with the shape of the ligand (**Fig. 6**). The electron density at these sites is distinct from apo STEP (**Fig. S10**), further arguing in favor of CoBrA being bound in our dataset. For Cys505, covalent binding necessitates a new side-chain rotamer, while the original, unbound rotamer remains present at partial occupancy. In the case of Cys518, covalent binding occurs at the original side-chain rotamer, selected from the pre-existing rotameric distribution of this residue in the apo STEP structure (**Fig. S10**).

**Figure 6.**
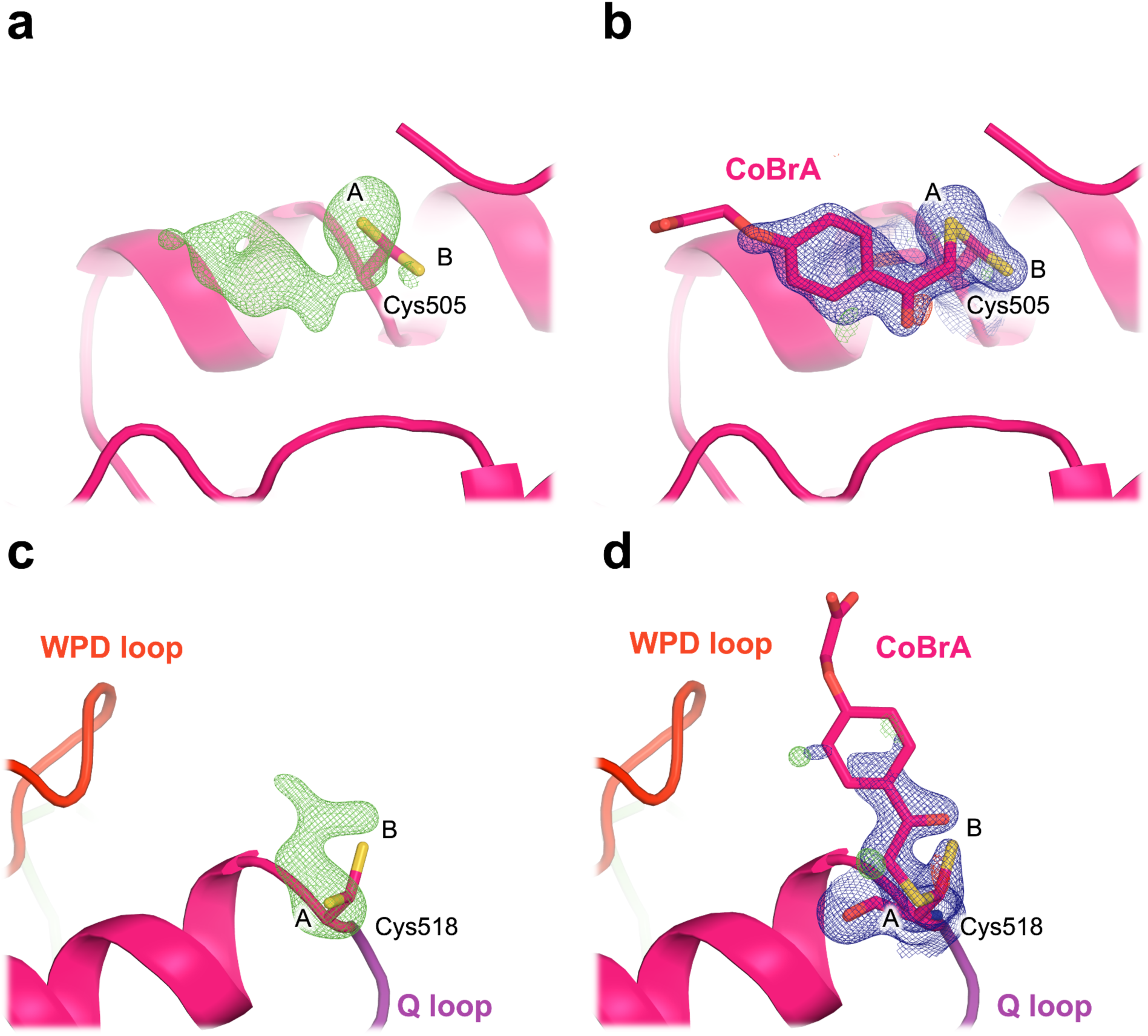
Covalent ligand binds to the distal Cys505 and Cys518 in STEP. a,c) Omit electron density (3.0 σ in green) for our structure soaked with CoBrA, showing unbiased evidence of covalent ligand binding. **b,d)** 2Fo-Fc (1.0 σ in blue) electron density for our refined structure with CoBrA modeled, showing good fit to density for the covalent ligand, although the flexible solvent-exposed end of the ligand is relatively disordered. Alternate conformations of the cysteine side chains are labeled as A and B. In **c-d)**, Cys518 is immediately adjacent in sequence to the catalytic Q loop (purple), with the catalytic WPD loop (red) also nearby.

In retrospect, there is evidence from other STEP structures that hints at the ligandability of Cys505. A crystal structure with an unmodified Cys505 (PDB ID 5ovx) ^11^ features an ordered Phe side chain which overlays with the central aromatic ring of CoBrA in our structure (**Fig. S11a**). Although this Phe is part of the recombinant purification tag, it nonetheless provides orthogonal validation of the physicochemical accessibility of this site. Furthermore, AlphaFold 2 ^39,40^ predicts with relatively high confidence (pLDDT > 85) that a linker region at the N-terminus of the catalytic domain in full-length STEP adopts an ordered helical conformation that can “dock” into the Cys505 binding site (**Fig. S11b**). This observation suggests that the Cys505 site may constitute a cryptic site, which would be significantly populated only upon ligand binding ^41^, in full-length STEP in human neurons.

Similar to the orthogonal evidence for Cys505, Cys518 is chemically modified (oxidized) in several previous STEP structures (PDB IDs 5ovx, 5ovr, 5ow1) (**Fig. S8h**, **Fig. S12**). This further corroborates the reactivity of Cys518, which supports what we observe in our structure.

To validate the covalent ligand binding events observed in our crystals, we used mass spectrometry in solution. The results showed strong labeling by CoBrA at several cysteines in wild-type STEP and both the C505S and C472S variants (**Fig. S13**, **Table S1**). Perhaps surprisingly given the presumed mechanism of action of CoBrA for other PTPs ^21^, the catalytic Cys472 was only minimally labeled.

Moreover, labeling was not detected at Cys505, arguing that the clear density for CoBrA at this residue in our crystal structure may be due to effects from the crystal lattice environment, or differences in labeling conditions (see Methods). Cys518, on the other hand, was among the most labeled residues, supporting our crystal structure. Cys488 was the most highly labeled residue overall, yet it is buried from solvent in all STEP crystal structures (**Fig. S14**). This suggests that local unfolding in this region likely occurs in solution to enable CoBrA binding. Notably, this area of the PTP catalytic domain fold (α4 helix) has allosteric properties in the STEP homolog PTP1B based on point mutations ^42,43^, suggesting a possible connection between conformational flexibility and allosterism.

We also tested whether CoBrA affects STEP enzymatic activity by binding at different cysteines. To do so, we generated C505S, C518S, and C505S/C518S mutants of STEP. For C505S, we were careful to use the same protein construct as was used for crystallography, to exclude the possibility that differences in the N-terminal purification tag region (**Fig. S11**) might contribute to differences in binding and/or activity. These mutations had weak to moderate effects on STEP activity (**Fig. S15**, **Table S2**) and did not alter inhibition of STEP by CoBrA (**Fig. S16**, **Table S3**). These observations suggest that CoBrA does not allosterically inhibit STEP by binding to Cys505 or Cys518 in solution, perhaps instead acting by promiscuous binding to multiple cysteine sites.

Altogether, although this particular covalent ligand does not allosterically modulate STEP activity, our results demonstrate that STEP contains several cysteines that are distal from the active site, chemically reactive, and covalently labeled by CoBrA in crystals and/or in solution. These sites thus represent promising starting points for development of new covalent ligands for STEP, including potential allosteric modulators or degraders.

## Discussion

This study underscores the value of X-ray crystallography under varied experimental conditions for revealing structural features of dynamic proteins. Here, we used different crystals of the protein tyrosine phosphatase STEP that (i) diffracted to particularly high resolution, (ii) were slowly dehydrated, and (iii) were soaked with a covalent ligand, with each dataset revealing new and unexpected structural insights.

In our high-resolution structure, we observed an ordered citrate molecule in the active site, supported by strong electron density (**Fig. 3**). Previous STEP structures have had electron density for competing molecules in the active site such as sulfates ^23^ or orthosteric inhibitors ^11^, which may have prevented binding of other molecules such as citrate in those structures. Given its high degree of coordination, our new citrate pose may be useful for structure-based drug design of small-molecule inhibitors of STEP. It sits at the structural nexus of several previously reported orthosteric inhibitors that extend in different directions from the catalytic pocket ^11^, but enjoys unique interactions with the adjacent E loop and WPD loop (**Fig. 3c**) that could potentially be exploited to modulate collective active-site dynamics of these two loops ^44^. Inhibitors that combine features of these different molecules could exploit the unique, atypically open (i.e. super-open) WPD loop conformation seen in all crystal structures of STEP thus far ^23,27^ to achieve specificity among the PTP family.

In our structure from a dehydrated crystal, the shifted conformation of the allosteric S loop was unanticipated. Yet this result is consistent with the notion that changes in protein crystal humidity ^45^ including those leading to dehydration ^46^ can significantly modulate protein conformational ensembles, adding to the toolkit of useful perturbations ^47^.

The new S loop conformation provides an unanticipated opening for structure-based design (**Fig. 4**, **Fig. S3**) to improve upon the existing allosteric activator scaffolds ^15^, dissect their somewhat mysterious mode of action, and/or convert them to allosteric inhibitors. The S loop is separated from the catalytic cysteine by only a single β-strand, and is immediately C-terminal of the α3 helix, which is involved in known allosteric mechanisms in the STEP paralog PTP1B ^48,49^. The S loop varies significantly among PTPs in terms of sequence as well as structure, including differences in loop length and conformation (**Fig. S4**). Despite this variability, multiple lines of evidence suggest the S loop is important for catalysis not only in STEP ^15^ but also in PTP1B: molecular dynamics (MD) simulations under different conditions show coupling of the S loop to the distal catalytic WPD loop ^50–52^, and mutation of a serine in the S loop prevents activation of PTP1B by protein kinase A, suggesting that the S loop contains a functionally relevant phosphorylation site ^53^. Notably, small-molecule fragment hits for PTP1B, derived both from computational reanalysis of crystallographic fragment screening data ^54^ and from MD simulations validated by crystallography ^55^, bind directly adjacent to the STEP allosteric activator pose ^54^; this suggests that additional chemical matter is likely compatible with this allosteric site in STEP (and/or PTP1B).

In the same dataset, we observe evidence for an intramolecular disulfide bond between the catalytic cysteine in the P loop and a backdoor cysteine in the nearby E loop. Such a disulfide in STEP had been previously observed using activity assays, mutagenesis, and mass spectrometry ^16^, but to our knowledge had not been structurally visualized in atomic detail before our study. The disordered density we observe immediately N-terminal to the backdoor Cys384 in the E loop, in contrast to the ordered state in other structures, suggests that disulfide formation may exploit the malleability of the E loop, which is evident from its variability among STEP structures ^23^. Our structure of an intramolecular disulfide for STEP complements those for other PTPs including LYP ^56^ and SHP2 ^31,57,58^ and for other phosphatases including PTEN ^59^ and Cdc25B ^60^. More generally, it adds atomistic detail to the growing picture of redox chemistry as a critical player in PTP regulation ^61–65^.

In our structure with CoBrA, we observe fortuitous covalent binding at Cys505 and Cys518, distal from the active site. This unexpected binding suggests that CoBrA is a promiscuous ligand, capable of labeling multiple cysteine residues beyond the active site. The lack of binding of CoBrA to Cys505 in solution, contrary to our crystal structure, may be due to a few possible factors. First, although there are no direct crystal contacts to this region in our crystal form, distributed effects from the crystal lattice could encourage labeling of Cys505 in crystals. Future studies could interrogate these effects in more detail, with varying protein constructs including full-length STEP (**Fig. S11**), different STEP crystal forms ^23^, and optimized compound variants. Second, conditions for LC-MS were chosen to minimize double labeling (see Methods). Given the apparent promiscuity of CoBrA, it is possible that with more permissive conditions, Cys505 would also be labeled.

The density we observe at Cys518 supports the modeling of CoBrA bound at this site (**Fig. 6**). This density, together with past structures of STEP with Cys518 modifications (**Fig. S8**), strongly suggests that Cys518 is chemically reactive. Additionally, Cys518 is substantially closer to the active-site WPD loop (∼7 Å) than is Cys505 (∼21 Å), so it has potential for covalent allosteric modulator discovery in the future as well.

Among human PTPs, Cys505 is nearly unique to STEP, as it is present in only the two closest homologs (HePTP and PTPRR) ^34^, and Cys518 is entirely unique to STEP — underscoring the potential of these residues for achieving specificity in allosteric modulation ^34^. Even if these sites are ultimately shown to be allosterically uncoupled from the active site, benign ligands could be developed into PROTAC degraders ^66^ with similar advantages in specificity.

Finally, the recent report of multiple non-orthosteric binders identified by high-throughput protein thermal shift assays ^67^ suggests that the map of ligandability for STEP is chiefly uncharted and broader than previously realized. Recently, crystallographic fragment screening ^68^ was employed quite fruitfully for the homolog PTP1B ^49,54,69^. This emerging technique is a promising near-future avenue toward mapping the ligandability of STEP more comprehensively, with an eye toward developing specific, potent, allosteric, small-molecule modulators for STEP to combat a spectrum of challenging neurological diseases.

## Methods

### Protein expression and purification

To obtain protein samples for crystallography, a plasmid containing the catalytic domain (residues 258–539) of STEP (PTPN5) with an N-terminal 6xHis & TEV cleavage site was obtained via Addgene from Nicola Burgess-Brown (Addgene plasmid #39166; http://n2t.net/addgene:39166; RRID:Addgene_39166). This was transformed into BL21(DE3) Rosetta2 (pRARE2) *E. coli* cells (MilliporeSigma). The sequence of the insert was independently verified using Sanger sequencing, with standard T7 promoter primers. Amino acid residue numbering in this paper follows UniProt isoform P54829-3 to ensure consistency with the consensus from almost all previously published STEP structures ^11,23,27^, the only past counterexample for human STEP being PDB ID 6h8r ^15^.

#### Protein expression

Throughout the entire process, the antibiotics chloramphenicol (Cam) and ampicillin (Amp) were employed to sustain selection at working concentrations of 30 μg/mL and 100 μg/mL, respectively. Previously transformed cells from glycerol stocks were plated on an LB-Agar + Amp + Cam plate and incubated overnight at 37°C. Individual colonies were selected and cultured overnight at 18°C in LB + Amp + Cam starter cultures (10 mL), shaking at 180 rpm.

Subsequently, this starter culture was added to baffled flasks containing 1 L of LB + Amp + Cam media and incubated until reaching an OD_600_ of 0.6–0.8 at 37°C, with shaking at 180 rpm. Expression was induced by adding isopropyl β-_D_-1-thiogalactopyranoside (IPTG) to a final concentration of 0.2 mM. Cultures were then incubated overnight at 18°C, shaking at 180 rpm, after which cells were harvested by centrifugation at 3000 rpm for 45 min, snap-frozen in liquid nitrogen, and stored at-80°C.

#### Protein purification

Frozen cellets (cell pellets) were thawed on ice, then 30 mL lysis buffer (50 mM HEPES pH 7.5, 500 mM NaCl, 5 mM imidazole, 5% v/v glycerol, 2 mM DTT) was added. One Pierce EDTA-free protease inhibitor mini-tablet per pellet was also added, followed by resuspension in a vortexer. Cells in the slurry were then lysed by 3 passages through a cell homogenizer (Avestin) operating with 1000 bar peak. Lysate was then centrifuged for 45 min at 50,000×g to spin down the cell fragments. The supernatant was filtered through a 0.22 μm filter to remove final cell debris.

A 5 mL Ni-NTA column (Cytiva) was equilibrated in freshly prepared low-imidazole buffer (50 mM HEPES pH 7.5, 500 mM NaCl, 30 mM imidazole, 5% v/v glycerol, 2 mM DTT). The lysate supernatant was applied to this column, washed with 2 column volumes (CV) of low-imidazole buffer, then gradient-eluted over 10 CV to 100% high-imidazole buffer (50 mM HEPES pH 7.5, 500 mM NaCl, 500 mM imidazole, 5% v/v glycerol, 2 mM DTT), collecting in 5 mL fractions. The STEP-containing fractions eluted around the 40% gradient mark were collected, concentrated using a 15 mL Centriprep 10 K spin-concentrator (Millipore) to a final volume of 5 mL, and filtered through a syringe-mount 0.22 μm filter to remove unidentified precipitate.

A Sephadex 20/200 column (Cytiva) was equilibrated with 2 CV of SEC buffer (50 mM HEPES pH 7.5, 500 mM NaCl, 5% v/v glycerol, 2 mM DTT). The concentrated, filtered Ni-binding fraction was injected onto a 5 mL loop, loaded onto the column, and fractionated over 2 CV, collecting 1 mL fractions. Two peaks were observed, and the fractions corresponding to the largest, STEP-containing peak were pooled.

A HiTrap Q HP anion-exchange column (Cytiva) was equilibrated with 2 CV of low-salt buffer (50 mM HEPES pH 7.5, 10 mM DTT). The pooled peak from size-exclusion chromatography was diluted to a final volume of 100 mL by addition of low-salt buffer, and filtered through a 0.22 μm bottle-top vacuum filter (Celltreat). This was then applied to the Q column, washed with 2 CV of low-salt buffer, and then gradient-eluted over 5 CV to 100% high-salt buffer (50 mM HEPES pH 7.5, 1000 mM NaCl, 10 mM DTT) collecting 5 mL fractions. A single STEP-containing peak was collected at 40% gradient mark.

This final STEP protein was concentrated in Centriprep 10 K spin-concentrators to 3 mL volume, and then further concentrated in Amicon 10 K spin-concentrators to a final concentration of 10 mg/mL, as measured by NanoDrop, and used fresh as the protein sample in crystallography. The identity of STEP vs. other proteins/contaminants was confirmed using SDS-PAGE gels at each step of the purification.

To obtain protein samples for activity assays with citrate, full-length human STEP_46_ was expressed and purified from *E. coli* as described previously ^67^.

To obtain protein samples for activity assays and mass spectrometry with CoBrA, we used the following steps:

#### PTP-encoding plasmid vectors

The pET vector encoding C-terminally 6xHis-tagged human STEP (pDK016; UniProtKB P54829, amino acids 282-565) was ordered from VectorBuilder ^70^. In this paper, we use CT-STEP as shorthand for this construct. Plasmids for the expression of all STEP mutants were generated by QuikChange site-directed mutagenesis, and desired mutations were confirmed via DNA sequencing by the Cornell Biotechnology Resource Center.

#### Protein expression and purification

BL21(DE3) *E. coli* cells containing the appropriate PTP-encoding plasmid were grown overnight at 37°C in LB. Cultures were diluted, grown to mid-log phase (OD_600_ = 0.5), induced with IPTG (1 mM), and shaken at room temperature overnight. The cells were harvested by centrifugation, resuspended and lysed with B-PER Bacterial Protein Extraction Reagent, and clarified by centrifugation. Enzyme purifications were carried out using HisPur Ni-NTA resin per the manufacturer’s instructions ^70^. Purified PTPs were exchanged into storage buffer (50 mM Tris pH 7.4, 0.5 mM EDTA, 0.01% Tween 20, 1 mM DTT), concentrated, flash-frozen with liquid nitrogen, and stored at-80°C. Protein concentration was determined using a NanoDrop spectrophotometer, and purity was assessed by SDS-PAGE.

### Crystallization

#### Citrate-bound STEP structure

For the citrate-bound STEP structure, precipitant well solution (30% PEG 3350, 120 mM Li_₃_[C_₆_H_₅_O_₇_] (lithium citrate), 100 mM bis-tris pH 5.65) was prepared fresh. A Mosquito (SPT Labtech) was used to prepare 96-well 3-drop Intelliplate Low-profile (Art Robbins Instruments) plates, in combination with hand-pipetting. 90 μL well solution was placed into the reservoir. One 4 μL drop was placed per well, using 1.75 μL of well solution pipetted by Mosquito, 0.5 μL of 10^-1^ diluted seed stock pipetted by Mosquito, and 1.75 μL of 10 mg/mL protein pipetted by hand. (The seed stock was stabilized in a Li_₂_SO_₄_-containing buffer.) Crystallization drops were incubated at room temperature. Crystals nucleated within 3 days, and grew over a week to around 80 x 80 x 20 μm. This crystal was looped and transferred to a 3 μL droplet containing a 1.2x solution of the mother liquor (30 mM HEPES pH 7.5, 120 mM NaCl, 60 mM bis-tris pH 5.5, 18% w/v PEG 3350, 60 mM Li_₃_[C_₆_H_₅_O_₇_] (lithium citrate)). 0.5 μL of 50% v/v DMSO was added to this 3 μL droplet. After approximately 10 minutes, the crystal was looped and cryocooled in liquid nitrogen.

#### Dehydrated STEP structure

For the dehydrated STEP structure, crystals were grown in Nextal EasyXtal 15-well hanging-drop trays. Precipitant well solution (0.2 M Li_2_SO_4_ (lithium sulfate), 0.1 M bis-tris pH 5.5, 30% PEG 3350) was prepared, and each reservoir was loaded with 400 μL. One 3 μL drop at a protein concentration of 10 mg/mL protein was hung per well, with 1:1 ratios of well solution to protein sample. Crystallization drops were incubated at room temperature. Crystals nucleated within 3 days, and grew over a week to around 140 × 70 × 40 μm. It is noteworthy that the crystal used for the dataset described here underwent an extended maturation period (∼6 months), leading to visually discernible dehydration of the crystallization drop. Before harvesting, the droplet containing the crystal was rehydrated using a 1.2x solution of the mother liquor (30 mM HEPES pH 7.5, 120 mM NaCl, 60 mM bis-tris pH 5.5, 18% w/v PEG 3350, 120 mM Li_2_SO_4_). The crystal was then looped, cryoprotected in LVCO (low-viscosity CryoOil, MiTeGen), and cryocooled in liquid nitrogen.

#### Covalently bound STEP structure

For the covalently bound STEP structure, the crystal was prepared using the same method as described above, using a 15-well hanging-drop tray. The crystal used for this dataset had dimensions of approximately 170 × 102 × 60 μm. In an effort to remove an anticipated active-site sulfate, the crystal was transferred to a 3 μL drop of a citrate-equivalent mother liquor, prepared to be equivalent ionic strength to the mother liquor: 25 mM HEPES pH 7.5, 100 mM NaCl, 50 mM bis-tris pH 5.5, 15% w/v PEG 3350, 50 mM Li_₃_[C_₆_H_₅_O_₇_] (lithium citrate). After 10 minutes, the soaking component 2-[4-(2-bromoacetyl)phenoxy]-acetic acid (CoBrA; C_10_H_9_BrO_4_; CAS number 29936-81-0; Cayman Chemical Company) was added to this pre-soak droplet, to achieve a final concentration of at least 10 times the concentration of the protein. Briefly, 0.5 μL of solubilized CoBrA (10.6 mM in 50% v/v DMSO/H_2_O) was added to this 3 μL droplet containing the crystal with a protein concentration of 5 mg/mL (approximately 0.1515 mM). CoBrA was allowed to react for approximately 10 minutes, then the crystal was looped and cryocooled in liquid nitrogen.

### X-ray diffraction

#### Citrate-bound STEP structure

For the citrate-bound STEP structure, X-ray diffraction data was acquired at the NYX beamline (19-ID) within Brookhaven National Laboratory’s NSLS-II synchrotron, with a cryostream operated at 100 K, using an X-ray beam energy of 12.658 keV and corresponding wavelength of 0.979 Å. The beam dimensions were set at 50 x 50 µm, with a flux ranging from 1.5–2.5×10^12^ ph/s. Exposures were held at 0.05 s over a 0.1° oscillation, with a total rotation of 360°, and no translation.

#### Dehydrated STEP structure

For the dehydrated STEP structure, X-ray diffraction data was acquired at the AMX beamline (17-ID-1) for Automated Macromolecular Crystallography at Brookhaven National Laboratory’s NSLS-II synchrotron, with a cryostream operated at 100 K, using an X-ray beam energy of 13.475 keV and corresponding wavelength of 0.920 Å. The beam dimensions were set at 7 x 5 µm, with a full flux of 4.25×10^12^ ph/s, and effective flux of 8.5×10^11^ ph/s with 20% beam transmission. Exposures were held at 0.01 s over a 0.1° oscillation, with a total rotation of 360°, and 125 μm translation.

#### Covalently bound STEP structure

For the covalently bound STEP structure, X-ray diffraction data was acquired at the FlexX beamline (ID7B2) for Macromolecular X-ray science at the Cornell High Energy Synchrotron Source (MacCHESS), with a cryostream operated at 100 K, using an X-ray beam energy of 11.3 keV and corresponding wavelength of 1.127 Å. The beam dimensions were set at 100 x 100 µm, with a flux of 8×10^11^ ph/s, and beam transmission of 100%. Exposures were held at 0.15 s over a 0.2° oscillation, with a total rotation of 360°, and no translation. Presumably due to the chemical changes involved with covalent binding of the bromo-compound, the crystal visibly changed color under irradiation.

### X-ray data processing and modeling

Data reduction and modeling were consistently applied across all three datasets using the DIALS data reduction pipeline ^71^. For all three datasets, resolution cutoffs were determined by taking into account CC_1/2_, I/sigma(I), R_merge_, and completeness.

Molecular replacement was initiated using DIMPLE ^72^ from CCP4 ^73^. Refinement of atomic coordinates, B-factors, and occupancies was performed with REFMAC ^74^ for initial stages, then with phenix.refine ^75^ for later stages. Manual adjustments to the models were made using Coot ^76^ between refinement rounds. Hydrogens were introduced using phenix.ready_set ^77^. The final rounds of refinement with phenix.refine involved the optimization of X-ray/stereochemistry weights and X-ray/ADP weights for 5 macro-cycles. In addition, given the high resolution of the citrate dataset, anisotropic B-factors were refined in the final stages, using riding hydrogens.

For the covalently bound STEP structure, two types of restraints files were used at each stage of refinement. First, the internal geometry of the ligand was enforced using a chemical restraints CIF file obtained from eLBOW ^78^. Second, the bond distance and angles for the covalent bonds between the ligand instances and the Cys side chains were specified in a manually curated restraints.edits file.

For both dehydrated and covalently bound STEP structures, we explored possible interpretations of the electron density for Cys505 and Cys518. First, in the covalently bound structure, the electron density for Cys505 is clearly consistent with CoBrA (**Fig. S10**). For Cys518, we attempted to model oxidative cysteine modifications previously observed in various structures in the PDB (CSO (S-hydroxycysteine), CSS (S-sulfinocysteine), CSU (S-sulfonylcysteine)), but these modifications fit the electron density poorly. We also considered a beta-mercaptoethanol (BME) adduct (CME(S-beta-mercaptoethylcysteine)), but it also failed to fit the electron density, and no BME was present in our experiments. We did not consider other cysteine variants involving post-translational modifications by enzymes (SCY, SCH). Second, in the dehydrated structure, we tried modeling the aforementioned modifications for both Cys505 and Cys518, but none fit the density satisfactorily. Due to the extended incubation period (∼6 months) prior to harvesting and data collection, we were unable to replicate this experiment in a reasonable timescale.

Solvent content was calculated using MATTPROB ^79–81^. To assess changes in the volume of the allosteric pocket, we used CASTp 3.0 ^82^ with default settings.

### Activity assays

For activity assays with citrate, STEP_46_ (one of the two major isoforms expressed in humans) phosphatase activity was measured at room temperature utilizing a standard 384-well plate format phosphatase fluorescence intensity assay using 3-O-methylfluorescein phosphate (OMFP) as the substrate ^83^. The reaction was performed in 25 µL 150 mM bis-tris (pH 6.5) buffer containing 50 mM NaCl, 0.5 mM EDTA, 0.01% Tween-20, 4 mM DTT. STEP_46_ concentration was 2.5 nM; OMFP was used at a concentration corresponding to its Michaelis-Menten constant for STEP_46_ (50 µM). Final concentrations for citric acid were 8, 4, 2, 1, 05, 0.25, 0.125, 0.0625, 0.0312, and 0.0156 mM. STEP and citric acid were incubated for 20 min at room temperature before the STEP reaction was started by the addition of a 5x OMFP working solution. Fluorescence intensity was measured for 10 min in kinetic mode using a Tecan Spark multimode microplate reader with an excitation wavelength of 485 nm, and an emission wavelength of 535 nm. The initial velocities were determined from the slopes of the linear progression curves of the STEP reaction. Rates were normalized using the no-enzyme and no-compound controls and analyzed using a nonlinear regression dose-response inhibition model (log inhibitor vs. response, variable slope, four parameters) and the program GraphPad Prism (version 10) to obtain the IC_50_ value.

For Michaelis-Menten activity assays and inhibition assays with CoBrA, phosphatase activity was measured by the rate of dephosphorylation of 6,8-difluoro-4-methylumbelliferyl phosphate (DiFMUP) as indicated by increasing emission at 440 nm. Reactions were pre-incubated for 30 minutes at 22°C and carried out in a 200 µL volume of PTP activity buffer (50 mM bis-tris at pH 6.5, 0.5 mM EDTA, 50 mM NaCl, 0.01% Tween 20, 1 mM DTT), enzyme, CoBrA or DMSO, and DiFMUP. Experiments were carried out at 0.1% v/v DMSO in **Fig. S15** and 1% v/v DMSO in **Fig. S16**. See **Fig. S15** for specific DiFMUP and enzyme concentrations and **Fig. S16** for specific DiFMUP, CoBrA, and enzyme concentrations. Enzyme-concentration uncertainty was estimated at 10% based on gel electrophoresis. Error bars on plotted data represent the standard deviation of triplicate measurements. Error bars in tabular data represent the standard error of the mean.

### Liquid chromatography tandem mass spectrometry

50 µL samples of 25 µM wild-type or mutant STEP were incubated with 125 µM CoBrA or DMSO in PTP storage buffer (50 mM tris at pH 7.4, 0.5 mM EDTA, 0.01% Tween 20, 1 mM DTT) for 30 minutes at 22°C. Samples were run on a NuPAGE^TM^ 4-12% bis-tris gel and stained with Coomassie brilliant blue G-250. The protein gel bands were then excised and subjected to in-gel trypsin digestion after reduction with dithiothreitol and alkylation with iodoacetamide. Labeled peptides were detected largely as described previously (detailed protocol in Supplemental Information) ^70,84,85^, and percent labeling for each cysteine in the STEP variants was calculated by dividing the sum of the total ion currents for cysteine-containing peptides labeled with CoBrA with the sum of the total ion currents for the corresponding unlabeled cysteine-containing peptides.

## Acknowledgements

We thank Sujit Chepuri for help with early experiments on the characterization of STEP inhibition by CoBrA.

Research reported in this publication was supported by the National Institutes of Health under Award Numbers R35GM133769 (to DK), RF1AG087011, R01AG065387, and R21AG067155 (to LT), R15GM071388 (to ACB), and by the NCI Cancer Center Support Grant P30CA030199. The content is solely the responsibility of the authors and does not necessarily represent the official views of the National Institutes of Health. We are also grateful for support from the Melvin and Phyllis McCardle Clause Scholarship program (fellowship to JW).

This work is based upon research conducted at National Synchrotron Light Source II (NSLS-II). NSLS-II is a United States Department of Energy (DOE) Office of Science user facility operated for the DOE Office of Science by Brookhaven National Laboratory under Contract DE-SC0012704. Use of the NYX beamline (19-ID) at NSLS-II is supported by the member institutions of the New York Structural Biology Center. Use of the AMX beamline (17-ID-1) at NSLS-II is supported by the Life Science Biomedical Technology Research resource, which is primarily supported by the NIH (NIGMS) through a Biomedical Technology Research Resource P41 grant (P41GM111244) and by the DOE Office of Biological and Environmental Research (KP1605010). We thank Kevin Battaile and the NYX staff for assistance with data collection at NYX, and Edwin Lazo and the AMX staff for assistance with data collection at AMX.

This work is also based upon research conducted at the Center for High Energy X-ray Sciences (CHEXS), which is supported by the National Science Foundation under award DMR-1829070, and the Macromolecular Diffraction at CHESS (MacCHESS) facility, which is supported by award 1-P30-GM124166-01A1 from the National Institute of General Medical Sciences, National Institutes of Health, and by New York State’s Empire State Development Corporation (NYSTAR). We thank Aaron Finke, David Schuller and the FlexX staff for assistance with data collection at the ID7B2 (FlexX) beamline at MacCHESS.

Liquid chromatography tandem mass experiments were conducted by the UMass Chan Medical School’s Mass Spectrometry Facility.

## Data Availability

The atomic coordinates and structure factor data for the three new STEP (PTPN5) crystal structures reported in this study have been deposited in the Protein Data Bank (PDB) under the following accession codes: 9EEX (citrate-bound STEP), 9EEY (dehydrated STEP), and 9EEZ (covalently bound STEP).

## Author Contributions

DAK and BTR conceived the project. BTR purified the protein and performed X-ray crystallography experiments. LG and AE performed structure modeling and refinement. SHK, JW, and YNH performed solution experiments including activity assays and mass spectrometry. LG, AE, and DAK wrote the manuscript with input from all authors. DAK supervised the structural biology aspects of the project. ACB and LT supervised solution experiments and edited the manuscript.

