## Supplemental Text, Figures, and Tables for "Three STEPs forward: A trio of unexpected structures of PTPN5"

### Supplemental Methods

#### Liquid chromatography tandem mass spectrometry

50 µL samples of 25 µM wild-type or mutant STEP were incubated with 125 µM CoBrA or DMSO in PTP storage buffer (50 mM tris at pH 7.4, 0.5 mM EDTA, 0.01% Tween 20, 1 mM DTT) for 30 minutes at 22°C. Samples were run on a NuPAGE™ 4-12% bis-tris gel and stained with Coomassie brilliant blue G-250. The protein gel bands were then excised subjected to in-gel trypsin digestion after reduction with dithiothreitol and alkylation with iodoacetamide. Peptides eluted from the gel were lyophilized and reconstituted in 20 µL of 5% acetonitrile containing 0.1% (v/v) formic acid (Suprapur, catalog no. 1116700250, EMD Millipore Corporation) and analyzed on a NanoAcquity Ultra Performance LC (UPLC) system (Waters Corporation, Milford, MA) coupled to a Orbitrap Fusion Lumos Tribrid mass spectrometer (Thermo Fisher Scientific Inc., Waltham, MA). A 2-µL injection was loaded at 4 µL/min for 4 min onto a custom-packed fused silica pre-column [Kasil frit, 100 µm internal diameter (I.D.)] with 2 cm of ProntoSIL C18AQ (Bischoff Chromatography, 200 Å, 5 µm). Peptides were then separated on a 75 µm I.D. fused silica analytical column packed with 25-cm Magic C18AQ (Bruker-Michrom, 100 Å, 3 µm) particles to a gravity-pulled tip. Peptides were eluted at 300 nL/min using a linear gradient from 5 to 35% of mobile phase B [0.1% (v/v) formic acid in acetonitrile], mobile phase A [0.1% (v/v) formic acid in water], in 115 min. Ions were introduced by positive electrospray ionization via liquid junction at 1.4 to 1.6 kV into a Orbitrap Fusion Lumos Tribrid mass spectrometer. Mass spectra were acquired from mass/charge ratio (m/z) 300 to 1750 at a resolution of 120,000 (m/z 200), maximum injection time of 50 ms using an AGC target of $4 \times 10^5$, and data-dependent acquisition (3-s cycle time) for tandem mass spectrometry by HCD fragmentation using an isolation width of 1.6 Da, max fill time of 22 ms, with an AGC target of $5 \times 10^4$. Peptides were fragmented using a collision energy of 30%, and fragment spectra were acquired at a resolution of 15000 (m/z 200). Raw data files were processed with Proteome Discoverer (Thermo Fisher Scientific, version 2.5) and searched with Mascot Server (Matrix Science, version 2.8) against the Human (Swiss-Prot) FASTA file (downloaded September 2023). Tryptic specificity (up to two missed cleavages), a 10-ppm mass tolerance for the precursor, and a 0.05-Da mass tolerance for the fragments were used as search parameters. Variable modifications of acetyl (protein N-term), pyro glutamic (N-term glutamine), oxidation (methionine), and CoBrA (cysteine) were selected. All nonfiltered search results were processed by Scaffold (Proteome Software Inc., version 5.3.0) with threshold values set at 95% for peptides and 99% for proteins (two peptide minimum) using the Trans-Proteomic Pipeline (Institute for Systems Biology). Search results were used to create spectral libraries for the Skyline software (University of Washington), which was used to quantitate selected peptides using precursor intensity data from extracted ion chromatograms. Percent labeling for each cysteine in the STEP variants was calculated by dividing the sum of the total ion currents for cysteine-containing peptides labeled with CoBrA with the sum of the total ion currents for the corresponding unlabeled cysteine-containing peptides.

### Supplemental Figures & Tables

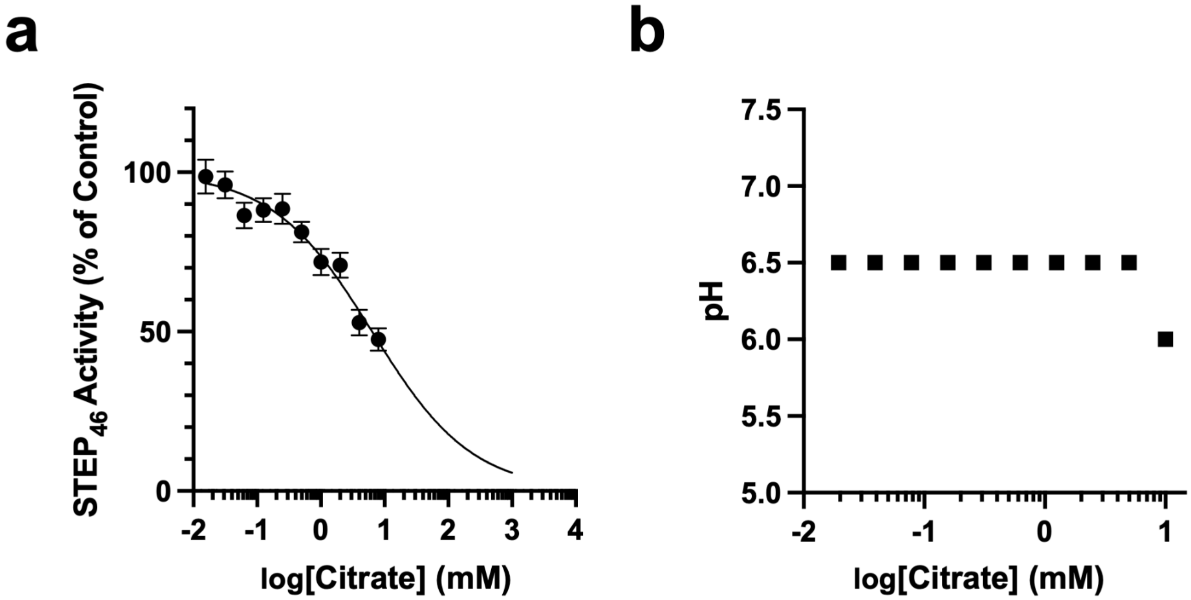


**Figure S1. Citrate weakly inhibits STEP activity.**
**a)** 3-O-methylfluorescein phosphate (OMFP) activity assay as a function of citrate concentration. $IC_{50}$ = 6.4 mM (95% CI: 5.2–8.2 mM).
**b)** pH of assay solution as a function of citrate concentration. Citrate concentrations above 10 mM were not included due to observed changes in pH.

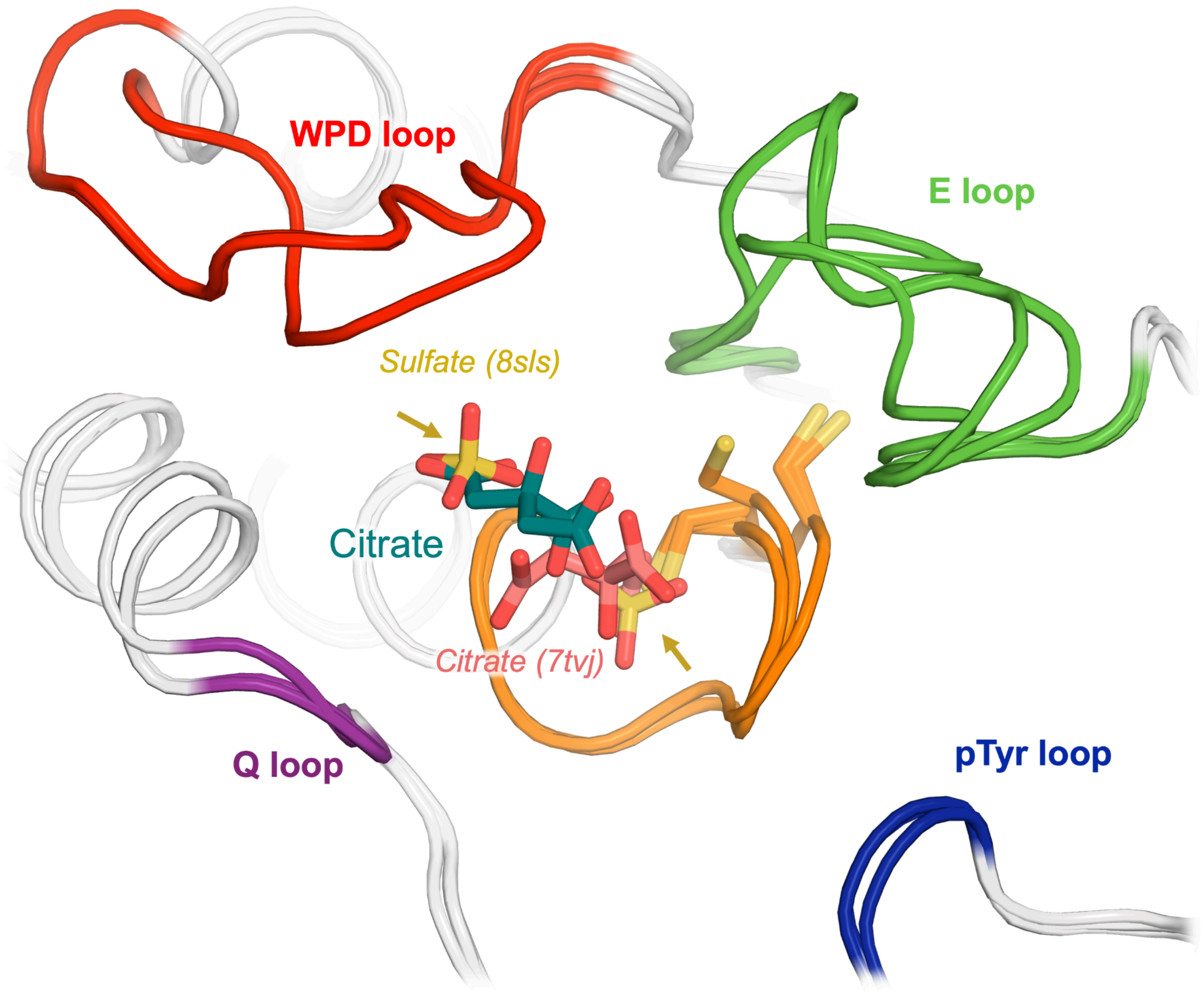


**Figure S2. Citrate or sulfate molecules bound to distinct active-site subsites in different PTP structures.** Alignment of three PTP structures, including our new citrate-bound structure, with citrate or sulfates bound in the active site: STEP (PDB ID 8sls) with two sulfates bound in the "top" and "bottom" subsites [23] (gold), our new structure of STEP with a citrate in the top subsite only (dark green), and SHP2 (PDB ID 7tvj) with a citrate in the bottom subsite only [31] (salmon).

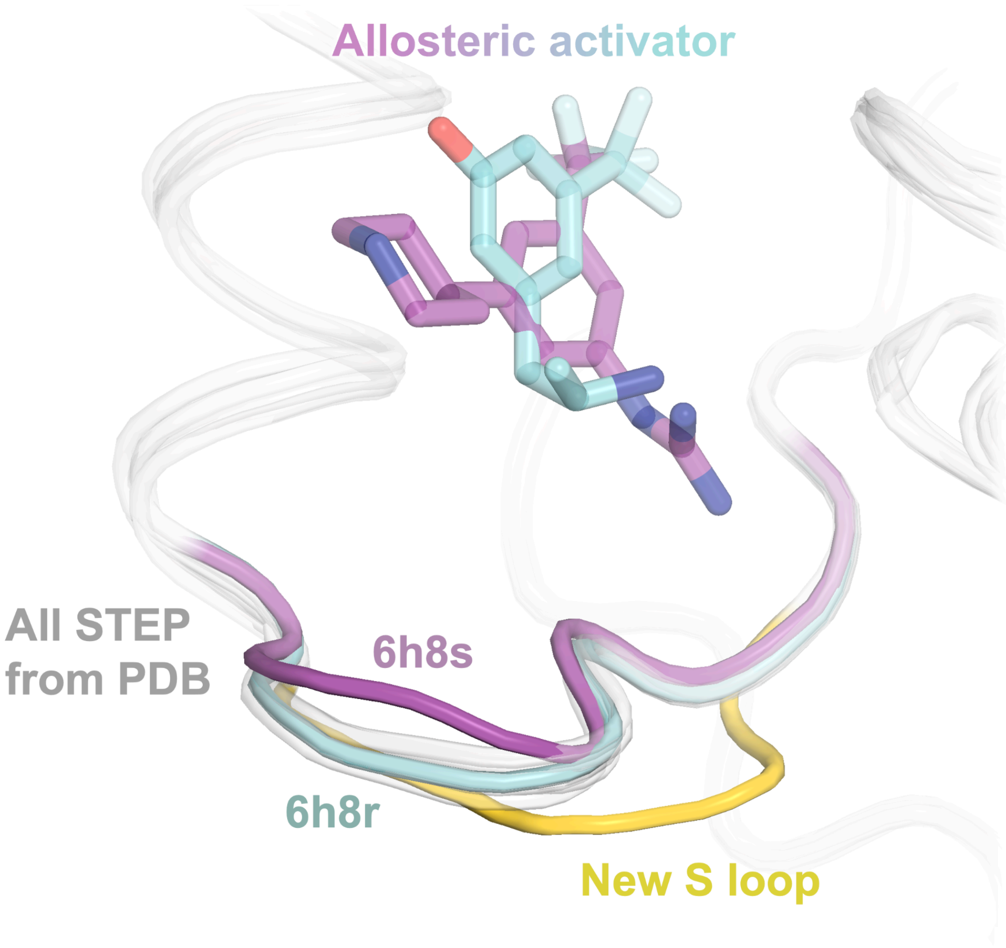


**Figure S3. New S loop conformation is different from all published STEP structures.**
The allosteric S loop in our dehydrated structure exhibits a distinct open conformation (gold), setting it apart from all the preceding STEP structures, including those with an S loop bound to an allosteric activator (cyan/purple) and those with no ligand bound in the allosteric site (transparent gray).

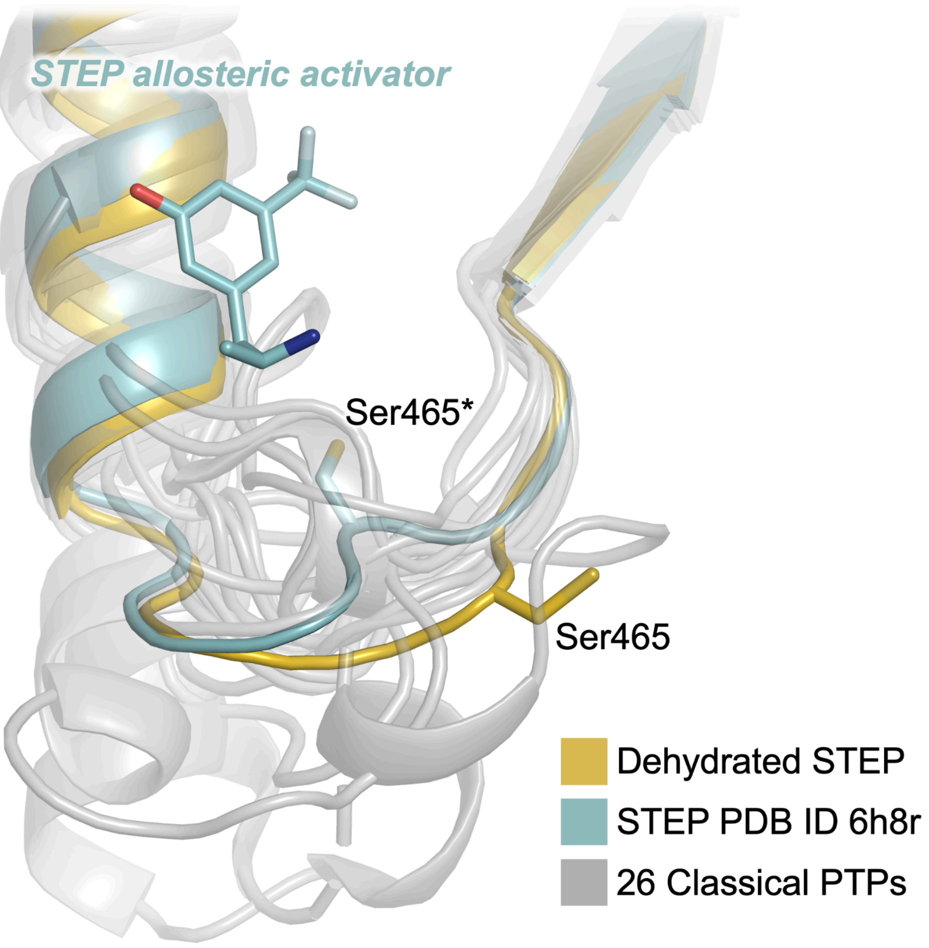


**Figure S4. S loop conformational diversity among structurally characterized classical PTPs.**
Alignment of representative crystal structures of 26 structurally characterized classical PTPs (out of the 37 total), including dehydrated STEP (gold) and STEP bound to an allosteric activator (PDB ID 6h8r, cyan, with the activator shown in sticks). The alignment highlights the conformational diversity of this region. Asterisks (*) indicate our renumbering of the non-standard residue numbering in 6h8r (see Methods).

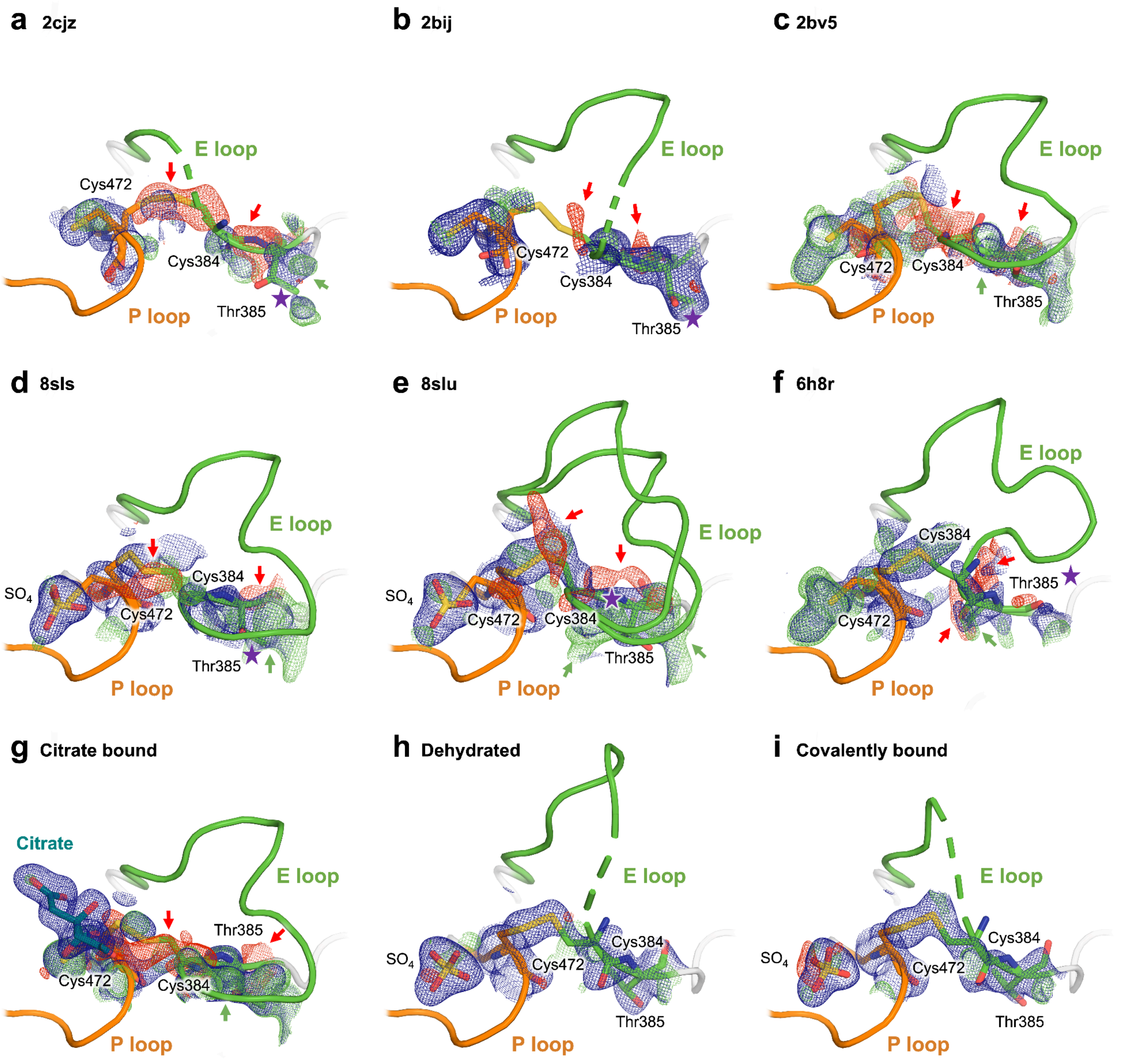


**Figure S5. Electron density evidence for Cys472-Cys384 disulfide in different STEP structures.** Several structures of STEP were remodeled and rerefined with a putative intramolecular disulfide bond between the catalytic Cys472 and the nearby Cys384 from the E loop. For each panel, refined 2Fo-Fc (1 σ contour, blue) and Fo-Fc (+/- 3 σ, green/red) electron density maps are shown. Ramachandran outliers are indicated by purple stars.

**a-g)** Negative Fo-Fc peaks, poor fit to 2Fo-Fc density, and/or Ramachandran outliers argue against the presence of a disulfide bridge between the catalytic and backdoor cysteines in numerous PDB structures of STEP, as well as our citrate structure. In **f)**, structure 6h8r was renumbered to match all other structures of STEP.

**h-i)** 2Fo-Fc density fits disulfide well, and Fo-Fc peaks are minimal. The E loop region preceding Cys384 appears disordered in both the dehydrated and covalently bound structures.

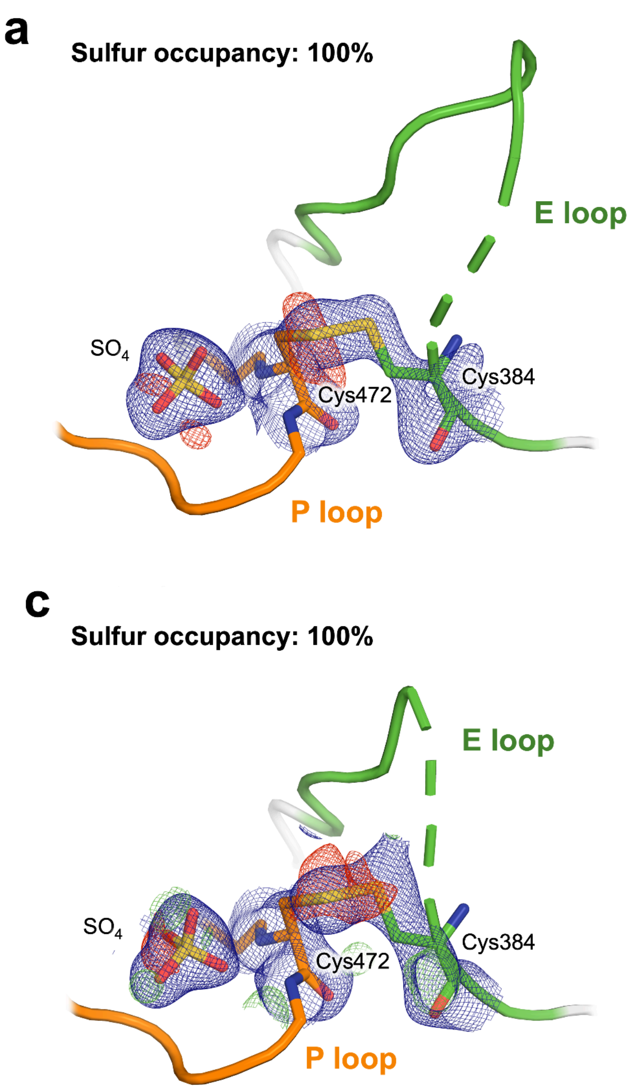


**Figure S6. Radiation-induced disruption of disulfide bonds leads to lower occupancy of sulfur atoms.**
**a)** Dehydrated STEP structure disulfide bond refined at full occupancy for both cysteine residues, including the sulfur atoms, shows negative density peaks (Fo-Fc, -3 σ, red) in the electron density map near the sulfur atoms.
**b)** Dehydrated STEP structure disulfide bond refined at partial occupancy for the sulfur atoms (sulfur occupancies after refinement: 69%). Negative density peaks are resolved.
**c)** Covalently bound STEP structure disulfide bond refined at full occupancy for both cysteine residues, including the sulfur atoms, shows negative density peaks (Fo-Fc, -3 σ, red) in the electron density map at the sulfur atoms.
**d)** Covalently bound STEP structure disulfide bond refined at partial occupancy for the sulfur atoms (sulfur occupancies after refinement: 70%). Negative density peaks are resolved.
2Fo-Fc density shown in all panels at 1 σ.

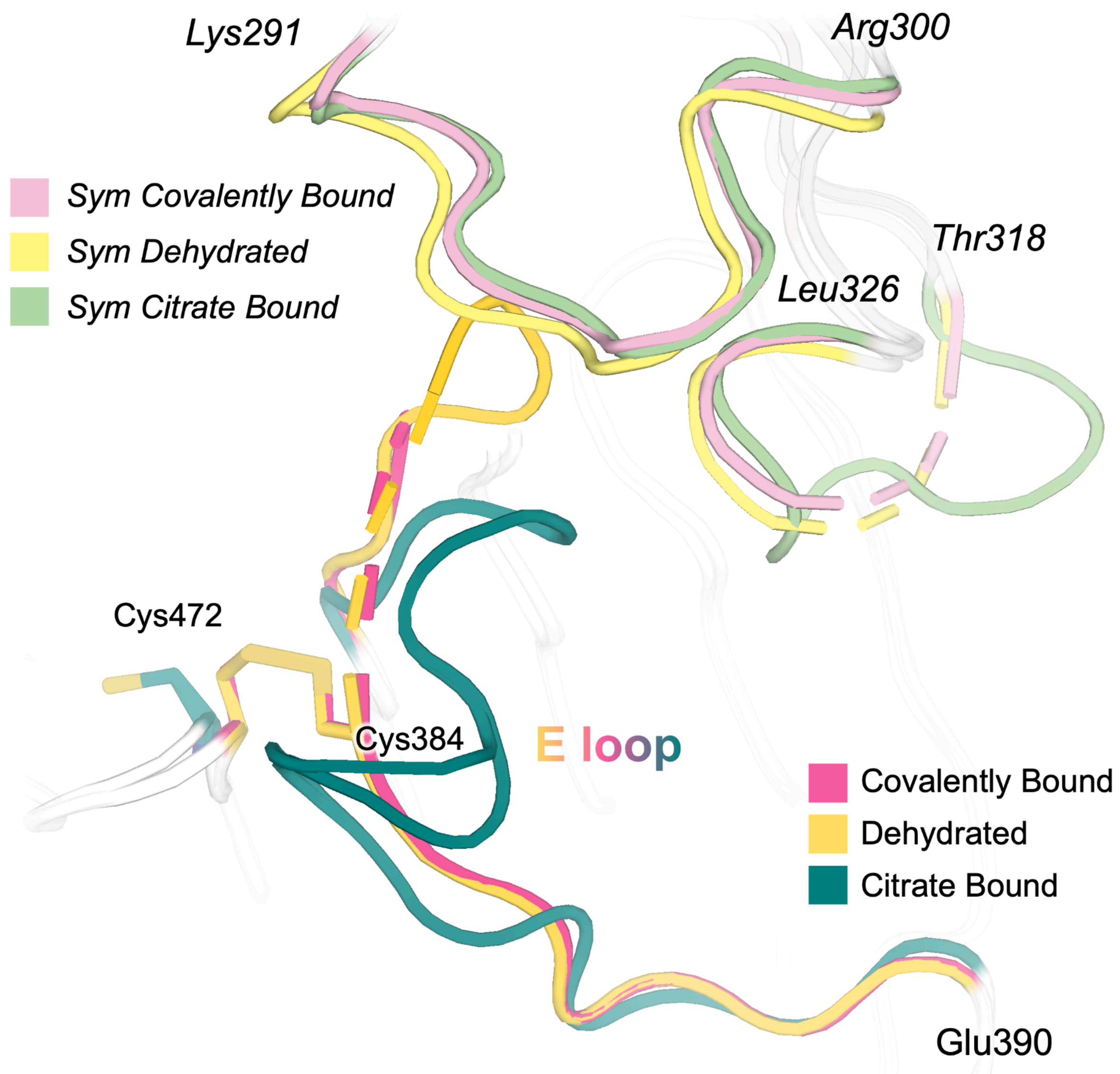


**Figure S7. E loop conformational change may propagate to nearby loops in the crystal lattice.** Alignment of the three STEP structures analyzed in this paper shows both the asymmetric unit (bottom, darker colors) and its symmetry mate (top, lighter colors). In the dehydrated and covalently bound STEP structures, a disulfide bond is observed between Cys472 and Cys384, while this bond is absent in the citrate-bound structure. In the structures with a disulfide bond, the E loop appears more dynamic or disordered. Two other loops, which are located nearby within the crystal lattice, also differ in conformation between our structures and/or show signs of relative disorder: the Thr318–Leu326 region and the Lys291–Arg300 region.

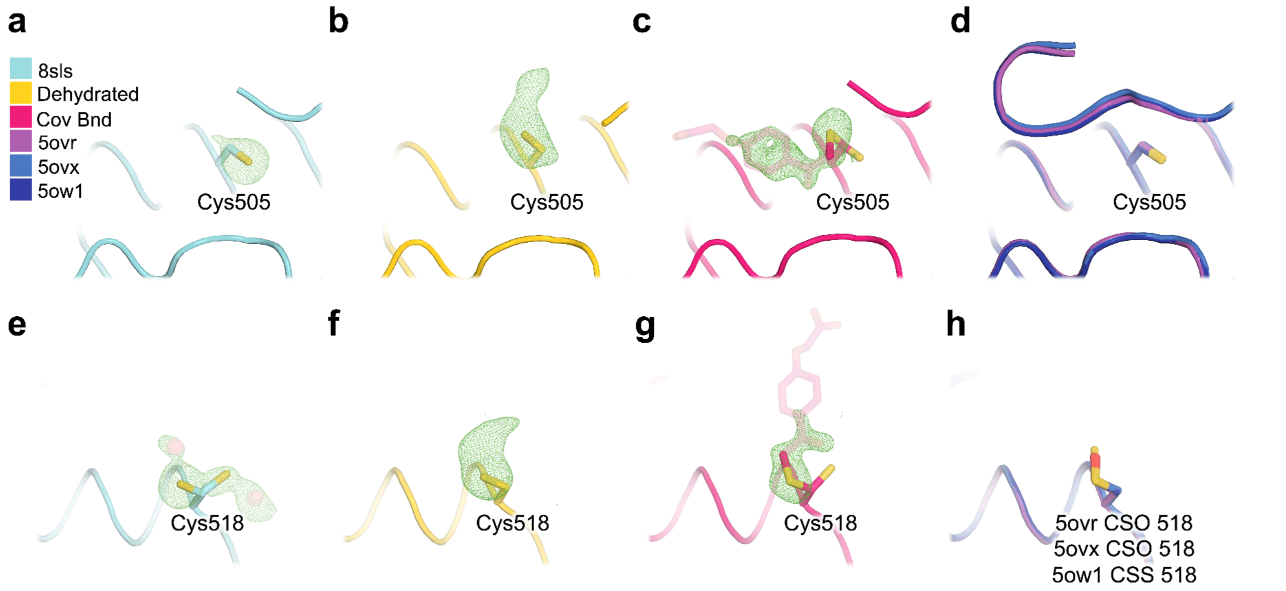


**Figure S8. Unbiased density for two non-catalytic cysteines reveals distinct chemical modifications in different structures.**

Two Cys sites, **a-d)** Cys505 and **e-h)** Cys518, are illustrated for several different STEP structures, as follows:

**a,e)** Fo-Fc (3.0 σ in green) omit map for an isomorphous apo structure (PDB ID 8sls) [23], showing no evidence for covalent ligand binding at either site.

**b,f)** Fo-Fc (3.0 σ in green) omit map of our dehydrated structure, revealing extra density at each site, suggesting covalent modification.

**c,g)** Fo-Fc (3.0 σ in green) omit map of our covalently bound structure, with the ligands modeled (transparent sticks), demonstrating a good fit in the density at both Cys505 and Cys518, supporting the presence of the covalent ligand at both sites.

**d,h)** Three other prior STEP structures (PDB IDs 5ow1, 5ovx, 5ovr) [11] showing distinct oxidized states of Cys518.

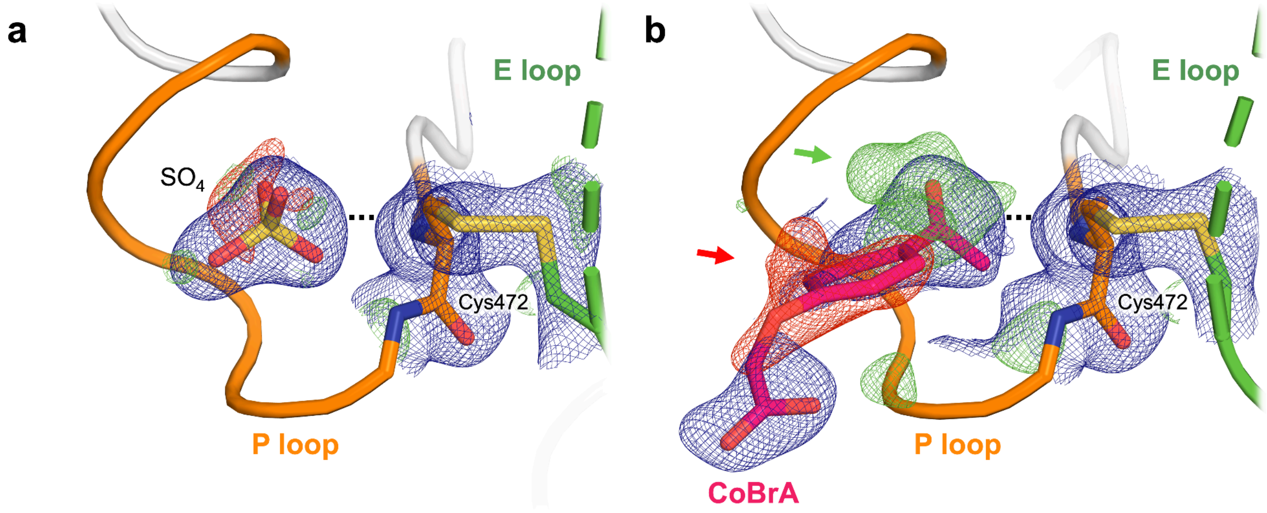


**Figure S9. Covalent ligand does not bind to the catalytic cysteine in STEP.**
**a)** Within the P loop, the catalytic cysteine Cys472 (participating in disulfide bond) is represented in sticks. The nearby electron density in the catalytic pocket is consistent with a bound sulfate, as seen in previous structures. 2Fo-Fc density contoured at 1.0 σ (blue); Fo-Fc density contoured at +/- 3.0 σ (green/red).
**b)** Putative model with the covalent ligand CoBrA positioned in the active site replacing the sulfate in **a)**. The refined 2Fo-Fc density map suggests a poor fit for CoBrA, including a persistent gap between the inhibitor and catalytic cysteine. The accompanying negative Fo-Fc density map features (see red arrow) further support this interpretation.

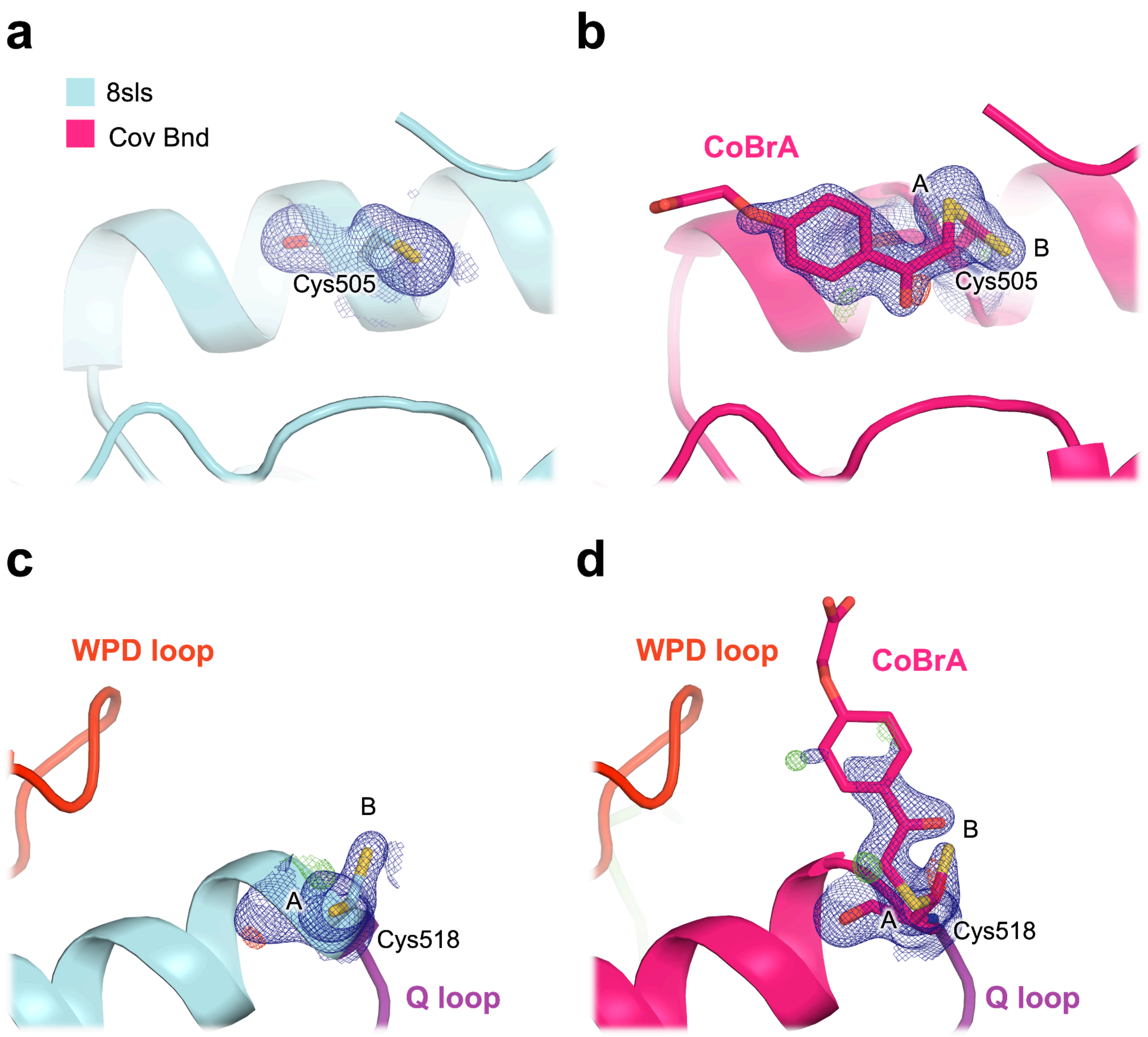


**Figure S10. Density for covalent ligand at Cys505 and Cys518 is distinct from apo density.**
**a,c)** 2Fo-Fc (1.0 σ in blue) electron density for an isomorphous apo structure (PDB ID 8sls) [23], showing no evidence for covalent ligand binding at either site.
**b,d)** 2Fo-Fc (1.0 σ in blue) electron density of our structure soaked with CoBrA with the ligand modeled, showing good fit in the density at both sites, although the flexible solvent-exposed end of the ligand is relatively disordered at Cys518.
Alternate conformations of the cysteine side chains are labeled as A and B.

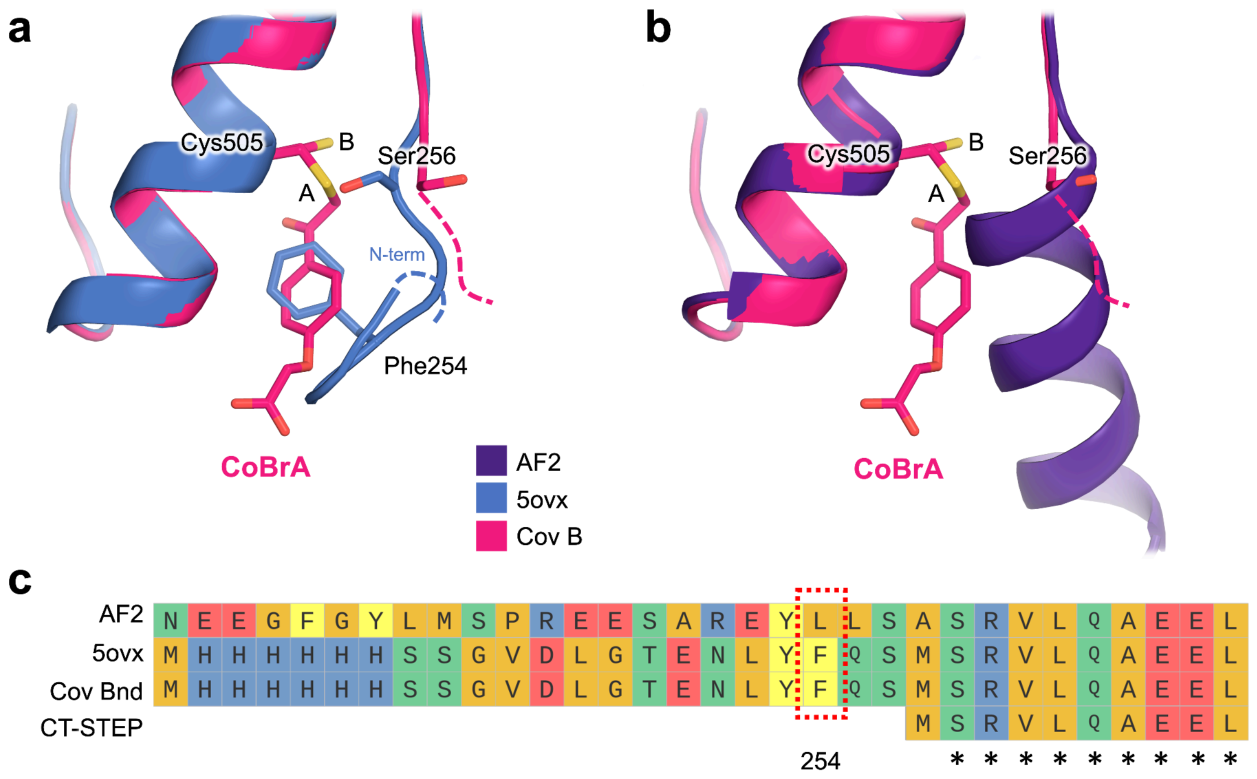


**Figure S11. Orthogonal evidence of ligandability and a potential cryptic site at Cys505.**

**a)** CoBrA covalently bound to Cys505 alternate conformation A in our structure of STEP (pink) partially mimics Phe254 (blue) of the N-terminal purification tag (TEV linker) in PDB ID 5ovx, which is disordered in our structure. Note that our structure and 5ovx have the same amino acid sequence including purification tag, but different crystal packing, which could help explain the difference in ordering at the N-terminal region of the catalytic domain.

**b)** Our structure with CoBrA overlaid with the AlphaFold 2 database [39,40] model for the full-length wild-type (WT) STEP sequence, showing that a helix may obstruct Cys505 to some degree in cells.

**c)** Sequence alignment of the N-terminal region for the AlphaFold model, 5ovx, our covalently bound structure, and a STEP construct with the purification tag located at the C-terminus (instead of the N-terminus), referred to here as CT-STEP. Dotted red box highlights Phe254 from the purification tag, which inserts into the pocket in 5ovx in panel **c)**. Asterisks (*) indicate that all amino acids are identical in the compared sequences.

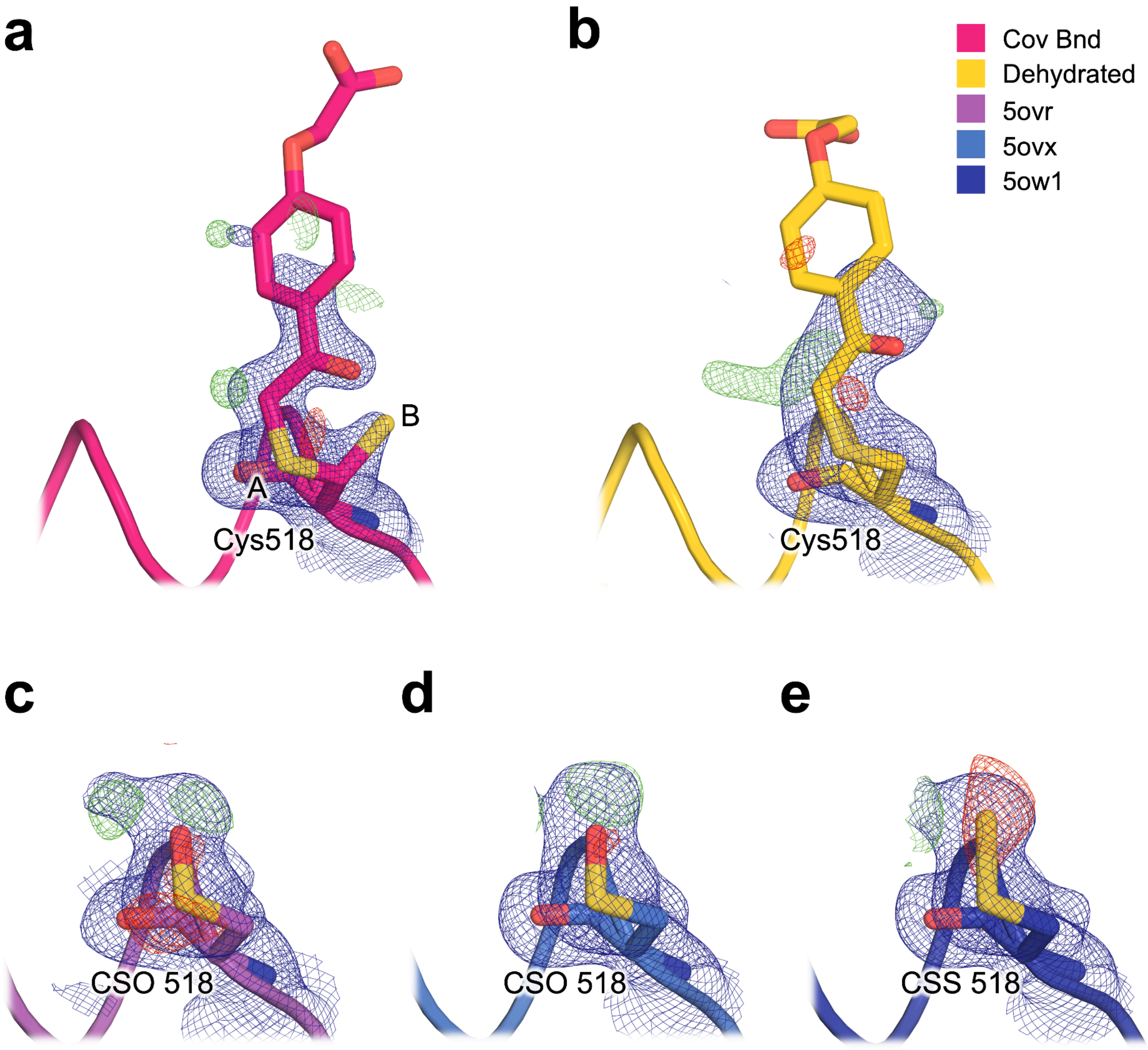


**Figure S12. Susceptibility to modification of Cys518.**
**a)** Covalently bound structure with CoBrA bound to Cys518.
**b)** Dehydrated STEP structure with hypothetical CoBrA modeled as bound to Cys518, despite its absence in the experiment, resulting in less convincing fit to density.
**c)** PDB ID 5ovr structure with S-hydroxycysteine (CSO) modification of Cys518.
**d)** PDB ID 5ovx structure with S-hydroxycysteine (CSO) modification of Cys518.
**e)** PDB ID 5ow1 structure with S-mercaptocysteine (CSS) modification of Cys518.
Each panel includes a refined 2Fo-Fc electron density map (1.0 σ in blue).

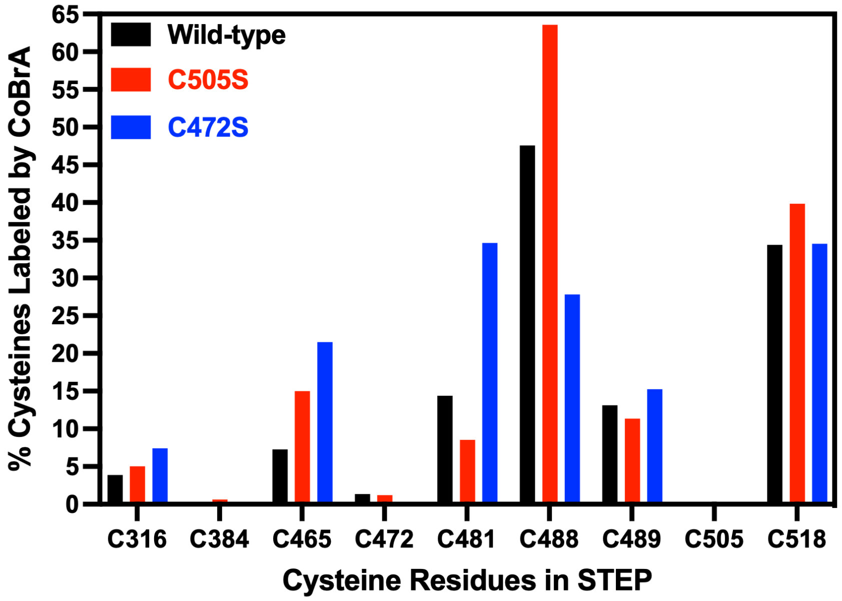


**Figure S13. LC-MS/MS shows covalent ligand binding to multiple cysteines in STEP.**
Percent labeling of cysteine residues in wild-type STEP and the C505S and C472S mutants by the covalent ligand CoBrA, detected by LC-MS/MS. 25 µM STEP was incubated with 125 µM CoBrA for 30 minutes at 22°C. This experiment uses the crystallography STEP construct. See **Table S1** for specific labeled peptides.

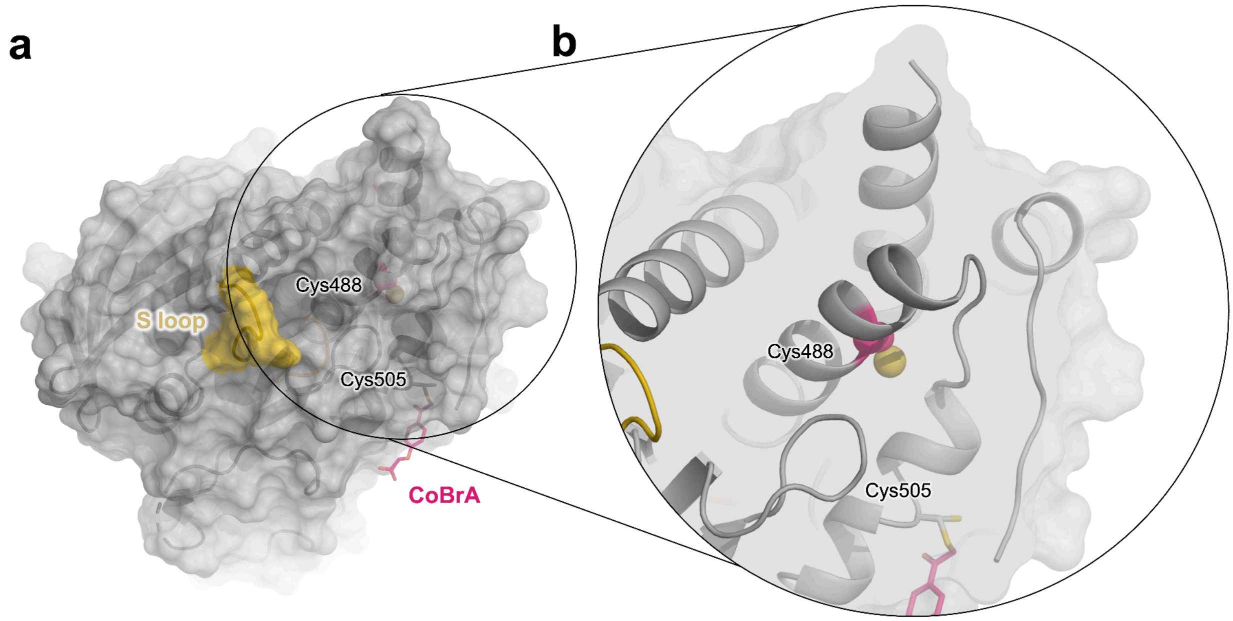


**Figure S14. Cys488 is not a surface-exposed cysteine.**
**a)** View of STEP distal to the active site showing the allosteric S loop (yellow) and CoBrA bound to Cys505 (pink sticks, right) for context. From this surface level view it can be observed that Cys488 (pink and yellow spheres, center-right) is buried, and may not be solvent accessible in this state.
**b)** Sliced zoom-in view of **a)**, showing that Cys488 (pink and yellow spheres, center) is buried and not exposed to the surface.

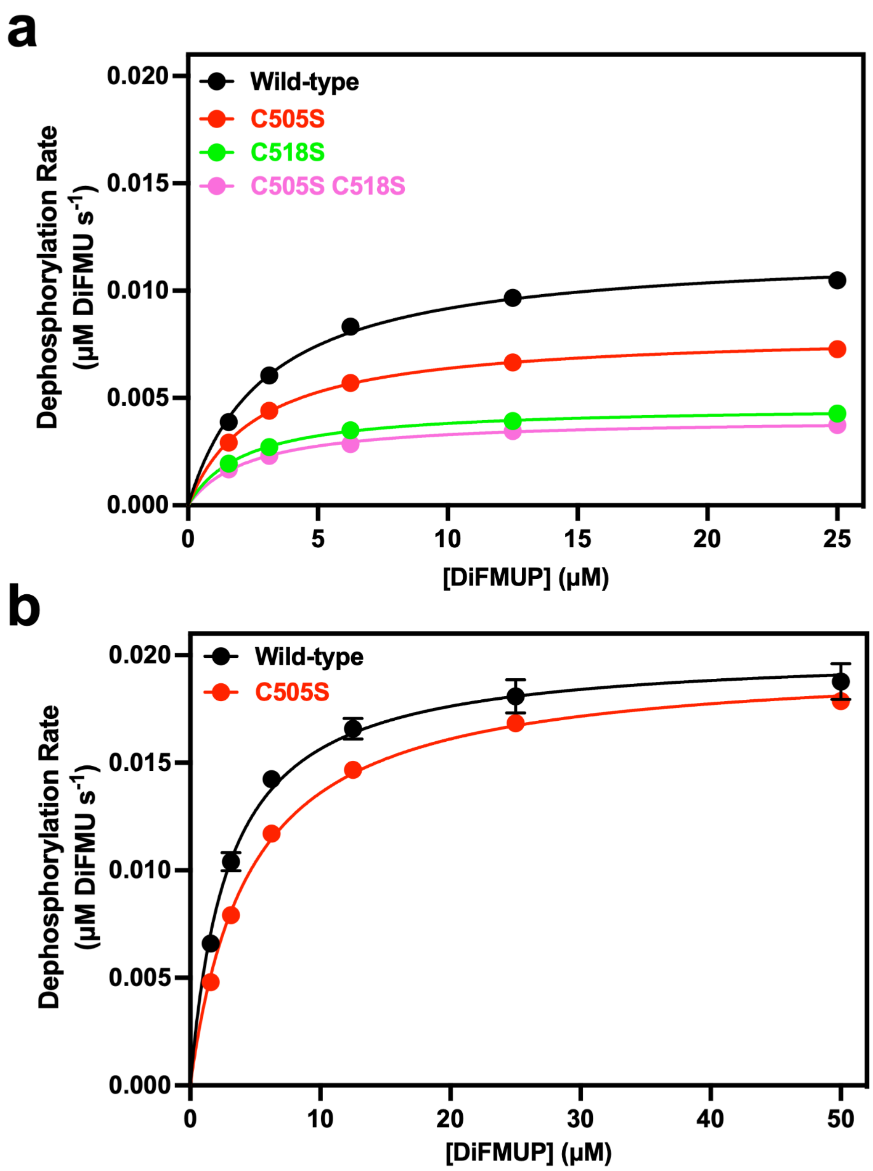


**Figure S15. Michaelis-Menten kinetics of STEP variants.**
**a)** Michaelis-Menten kinetics of STEP (C-terminally tagged, CT-STEP): wild-type, C505S, C518S, and C505S/C518S mutants.
**b)** Michaelis-Menten kinetics of STEP (N-terminally tagged, crystallography construct): wild-type and C505S mutant.
For each panel, PTP basal activities were assayed with the indicated concentrations of DiFMUP, and enzyme concentrations were 20 nM for all variants. See **Table S2** for kinetics constants. Error bars that are not visible are obscured by the plotted data.

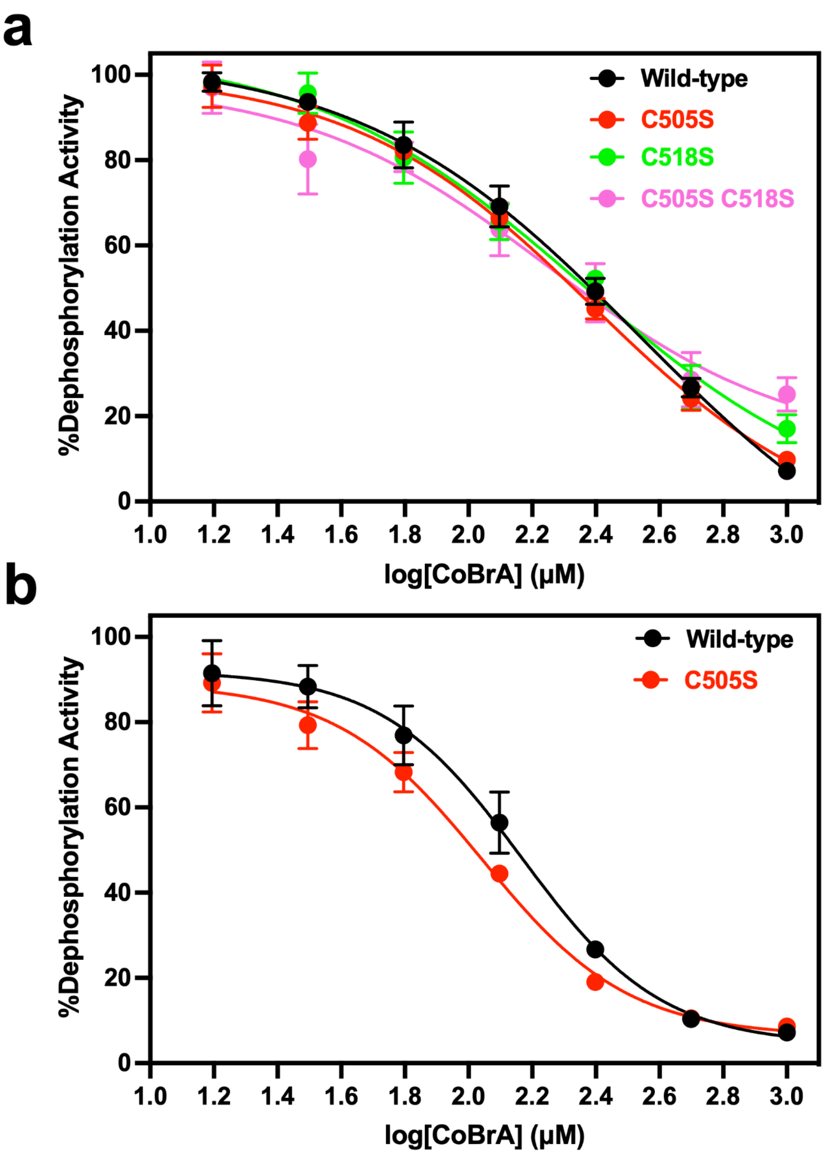


**Figure S16. Dose-dependent inhibition of STEP variants by covalent ligand.**
**a)** Dose-dependent inhibition of STEP (C-terminally tagged, CT-STEP): wild-type, C505S, C518S, and C505S/C518S mutants.
**b)** Dose-dependent inhibition of STEP (N-terminally tagged, crystallography construct): wild-type and C505S mutant.
For each panel, STEP variants (20 nM) were incubated with the indicated concentrations of CoBrA for 30 min at 22°C and assayed with 3 μM DiFMUP. See **Table S3** for $IC_{50}$ values. Error bars that are not visible are obscured by the plotted data.

| **Cysteine Residue** | **Predominant Peptide** | **Total Ion Current (TIC)** | **TIC, Labeled with CoBrA** | **TIC, Unlabeled** | **% Labeled with CoBrA** |
|---|---|---|---|---|---|
| 316 | VCLTSPDPDDPLSSYINANYIR | 1.25E+07 | 4.88E+05 | 1.20E+07 | 3.89 |
| 384 | CTEYWPEEQVAYDGVEITVQK | 6.14E+06 | 0 | 6.14E+06 | 0 |
| 465 | EVEEAAQQEGPHCAPIIVHCSAGIGR | 3.37E+07 | 2.45E+06 | 3.13E+07 | 7.26 |
| 472 | EVEEAAQQEGPHCAPIIVHCSAGIGR | 3.37E+07 | 4.57E+05 | 3.32E+07 | 1.36 |
| 481 | TGCFIATSICCQQLR | 2.23E+06 | 3.20E+05 | 1.91E+06 | 14.38 |
| 488 | TGCFIATSICCQQLR | 2.23E+06 | 1.06E+06 | 1.17E+06 | 47.58 |
| 489 | TGCFIATSICCQQLR | 2.23E+06 | 2.92E+05 | 1.94E+06 | 13.12 |
| 505 | QEGVVDILKTTCQLR | 4.20E+05 | 0 | 4.20E+05 | 0 |
| 518 | GGMIQTCEQYQFVHHVMSLYEK | 1.65E+07 | 5.68E+06 | 1.08E+07 | 34.40 |

**Table S1. Cysteine-containing peptides labeled by CoBrA from LC-MS/MS.**
Cysteine-containing peptides labeled by CoBrA in wild-type STEP. 25 μM STEP was incubated with 125 μM CoBrA for 30 minutes at 22°C, and the degree of labeling at indicated cysteine residues (red C letters) was determined by LC-MS/MS. This experiment uses the crystallography STEP construct.

| Construct | STEP Variant | $V_{max}$ (μM $s^{-1}$) | $K_M$ (μM) | $k_{cat}$ ($s^{-1}$) | $k_{cat}/K_M$ ($s^{-1}$ $mM^{-1}$) |
|---|---|---|---|---|---|
| ***CT-STEP*** | **Wild-type** | 0.0120 ± 0.0002 | 3.0 ± 0.1 | 0.60 ± 0.06 | 199 ± 6 |
| | **C505S** | 0.0080 ± 0.0001 | 2.7 ± 0.1 | 0.40 ± 0.04 | 152 ± 6 |
| | **C518S** | 0.0050 ± 0.0001 | 2.2 ± 0.1 | 0.23 ± 0.02 | 107 ± 4 |
| | **C505S/C518S** | 0.0040 ± 0.0001 | 2.4 ± 0.2 | 0.20 ± 0.02 | 86 ± 5 |
| ***NT-STEP*** | **Wild-type** | 0.0202 ± 0.0002 | 2.9 ± 0.1 | 1.0 ± 0.1 | 350 ± 10 |
| | **C505S** | 0.0198 ± 0.0001 | 4.5 ± 0.1 | 1.0 ± 0.1 | 218 ± 4 |

**Table S2. Michaelis-Menten kinetics constants for STEP variants.**
Michaelis-Menten kinetics constants for STEP variants assayed using DiFMUP.
First four data rows: C-terminally tagged, CT-STEP.
Last two data rows: N-terminally tagged, crystallography construct.

| Construct | STEP Variant | $IC_{50}$ (μM)* |
|---|---|---|
| ***CT-STEP*** | **Wild-type** | 230 ± 10 |
| | **C505S** | 210 ± 10 |
| | **C518S** | 210 ± 20 |
| | **C505S C518S** | 230 ± 40 |
| ***NT-STEP*** | **Wild-type** | 150 ± 10 |
| | **C505S** | 115 ± 8 |

**Table S3. $IC_{50}$ values for inhibition of STEP variants by covalent ligand.**
Half-maximal inhibitory concentrations ($IC_{50}$) of STEP variants with the covalent ligand CoBrA.
First four data rows: C-terminally tagged, CT-STEP.
Last two data rows: N-terminally tagged, crystallography construct.
*$IC_{50}$ values for covalent inhibition are generally dependent on the time and temperature of enzyme/inhibitor pre-incubation. The reported values were measured after 30-minute pre-incubations at 22°C.
